# Spectral Adaptive Repositioning for Flow-Based Single-Cell Perturbation Modeling

**DOI:** 10.64898/2026.11.02.763678

**Authors:** Shourya Verma, Mengbo Wang, Simran Kadadi, Nadia Atallah Lanman, Shahin Mohammadi, Ananth Grama

**Affiliations:** Computer Science; Comparative Pathobiology; Institute for Cancer Research, Purdue University; Broad Institute; Cytognosis Foundation

## Abstract

Predicting transcriptome-wide cellular responses to genetic and biomolecular perturbations is central to functional genomics and therapeutic target discovery. Existing methods assign gene positional encodings derived from arbitrary orderings or static co-expression masks built from unperturbed cells, and cannot capture the functional rewiring induced by perturbations. We present **SPEAR**, a generative framework built on two complementary positional mechanisms. Spectral positional encoding grounds each gene’s position in the geometry of the co-expression network via continuous coordinates derived from the graph Laplacian. Adaptive repositioning then derives each gene’s position from its perturbation-conditioned hidden state at every layer, allowing the attention genes exert on one another to reorganize without explicit supervision. Evaluated on genetic, pharmacological, and cytokine perturbation benchmarks, SPEAR demonstrates excellent recovery of differential gene-expression, distributional structure, and perturbation-specific identifiability. SPEAR also captures latent positional relationships that closely map to known transcription factor-target interactions. https://github.com/shouryaverma/SPEAR

## 1 Introduction

Predicting transcriptomic responses to cellular perturbations is a fundamental problem in functional genomics and target discovery. While high-throughput CRISPR screens and multiplexed pharmacological profiling have enabled the measurement of these responses across hundreds of conditions [1], the combinatorial space of possible genetic knockouts and chemical interventions grows exponentially with the number of targets [2]. This renders exhaustive experimental coverage practically infeasible. Machine learning models that generalize from observed perturbations to unseen combinations are essential, particularly for resolving multi-gene interactions, where epistatic or non-additive effects cannot be inferred from individual knockouts alone [3].

Recent generative frameworks address this challenge by formulating perturbation prediction as a distribution-to-distribution mapping, predominantly relying on Transformer-based architectures [4]. Despite this progress, a fundamental architectural limitation remains: Transformers require a notion of position to distinguish tokens, yet no biologically coherent positional encoding exists for genes. While genes possess a 1D sequence on chromosomes, sequential positional encodings fail to capture the complex, non-linear topology of gene regulatory networks; conversely, discarding positional encodings treats all genes as exchangeable, stripping the attention mechanism of any regulatory topology. Static binary attention masks attempt to compensate by restricting gene-level attention to co-expressed neighbors. However, these encode network structure as rigid on-off constraints rather than a continuous geometric prior. Because they remain identical across all cells and conditions, they cannot generalize beyond observed edges or capture the functional rewiring induced by a perturbation. We address the problem for cellular perturbation modeling, where tokens have no natural order, relationships are graph-structured, and those relationships rewire as a function of context.

We present **SPE**ctral **A**daptive **R**epositioning (**SPEAR**), which replaces sequential positional encodings and binary attention masks with two complementary mechanisms grounded in the continuous geometry of gene co-expression. Spectral co-expression encoding embeds each gene into a continuous coordinate space derived from the co-expression graph Laplacian. In this embedding, genes with similar co-expression patterns attend to one another with higher affinity, and each attention head can specialize to a different scale of co-expression community structure. Context-adaptive repositioning derives a scalar position for each gene at every backbone layer from its perturbation-conditioned hidden state. This allows the attention geometry to dynamically reorganize and reflect perturbation-induced functional rewiring, without requiring explicit supervision on gene-gene relationships. Both components are summed into a single rotation angle, so each head jointly encodes static co-expression proximity and perturbation-conditioned positional displacement.

Our contributions are: **1. Spectral co-expression encoding:** Continuous positional coordinates derived from the co-expression graph Laplacian and embedded into attention rotation angles via per-head learnable frequency projections, providing a topologically grounded geometric prior whose benefit is not reproduced by permuted or freely learned coordinates of the same dimensionality. **2. Adaptive repositioning:** Dynamic per-gene scalar positions derived from perturbation-conditioned hidden states at every backbone layer, allowing each gene’s influence in attention to adapt to specific perturbation contexts without explicit supervision. **3. State-of-the-art performance:** Robust predictive recovery across combinatorial genetic, pharmacological, and cytokine perturbation benchmarks.

## 2 Related Work

### Perturbation response prediction

Early deep learning approaches frame perturbation prediction as conditional generation in a learned latent space. scGen [5] models perturbation as a vector shift in a VAE, while CPA [6] disentangles cell state, perturbation identity, and covariates to support combinatorial drug and genetic knockouts. Both treat perturbation effects as population averages and ignore co-expression structure among genes. GEARS [7] addresses out-of-distribution generalization by routing gene embeddings through a GNN over a co-expression and Gene Ontology knowledge graph, enabling prediction for gene combinations never individually perturbed. Foundation models trained on tens of millions of single-cell transcriptomes, Geneformer [8] and scGPT [9] learn transferable gene representations but assign positions by expression rank or integer index, discarding co-expression topology. More recent work shifts to distribution-level objectives: STATE [10] trains a large transformer on over 100 million perturbed cells with a kernel-based population alignment loss, and CellFlow [11] applies flow matching in a low-dimensional PCA space to learn perturbation-induced cell state transitions. scDFM [4] applies conditional flow matching directly in gene expression space with a differential transformer backbone and a static binary co-expression attention mask. No prior perturbation model encodes co-expression topology as a continuous geometric prior or allows the influence genes exert on one another in attention to adapt dynamically to perturbation context.

### Positional encodings for transformers

Rotary Position Embedding (RoPE) [12] encodes relative position by rotating query and key vectors before the attention dot product so that attention scores depend only on the pairwise positional distance. This has become the standard rotary mechanism for continuous relative encoding. Standard RoPE requires integer token indices and is inapplicable to gene sets without a biologically meaningful ordering. In the graph learning literature, positional encodings derived from the eigenvectors of the graph Laplacian have been used to assign continuous structural coordinates to nodes in graph transformers [13, 14, 15], encoding topological proximity as a geometric prior rather than a discrete adjacency constraint. Context-dependent encodings that derive positions from token representations rather than fixed precomputed indices have also been explored for sequence length generalization in language models [16, 17]. Graph Laplacian encodings have been studied for static topology, and context-dependent positions for sequence-length generalization; neither has been applied to encoding perturbation-conditioned gene co-expression structure in single-cell modeling.

## 3 Methods

### Problem Formulation

Let *G* = {*g*_1_, …, *g*_*G*_} denote a fixed vocabulary of *G* genes. A cell is represented as an expression vector **c**∈ ℝ^*G*^, where *c*_*i*_≥ 0 is the normalized count of gene *g*_*i*_. A perturbation *ρ* is a set of one or more knocked-out gene identifiers. Given a source distribution *p*_0_ of unperturbed control cells and a target distribution *p*_1_ of cells under perturbation *ρ*, the task is to learn a generative model *T*_*ρ*_ : *p*_0_ →*p*_1_ that transfers control cells to their perturbed counterparts and generalizes to held-out *ρ*. We frame this as flow matching: a velocity field *v*_*θ*_(**x**_*t*_, *t*, **c**_ctrl_, *ρ*) is trained to transport samples from a noise distribution *p*_noise_ to *p*_1_, conditioned on the paired control cell **c**_ctrl_ and perturbation *ρ*. Integrating the learned ODE from *t* = 0 to *t* = 1 at inference produces a sample from the predicted perturbed distribution. Each gene *g*_*i*_ is assigned a token index *τ*_*i*_ ∈ {0, …, |*V*|−1} via a vocabulary *V* constructed from highly variable genes. A batch operating on gene subset *S* ⊆ *G* of size *N* is represented as ***τ*** ∈ ℤ ^*B*×*N*^, where *B* is the batch size.

### Spectral Co-expression (SPE)

Integer positional encodings are semantically meaningless for genes, since they have no natural order. We instead derive continuous positional coordinates from the topology of the gene co-expression graph [18], precomputed once from training data. Let **X**∈ ℝ^*C*×*G*^ be the log-normalized training expression matrix. We column-normalize **X** to zero mean and unit *l*_2_ norm to obtain 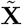, then form the symmetric co-expression adjacency: 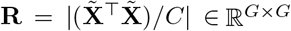, *R*_*ii*_ = 0, where the element-wise absolute value treats co- and anti-correlated gene pairs symmetrically. To focus on the strongest co-expression relationships and suppress long-range noise, each row of **R** is sparsified to its top-*k* entries: *A*_*ij*_ = *R*_*ij*_ **1**[*j* ∈ *N*_*k*_(*i*)], **A** ← max(**A, A**^⊤^), where *N*_*k*_(*i*) is the index set of the *k* genes with largest *R*_*ij*_, and the max symmetrizesthe result. The symmetric normalized Laplacian is then **L** = **I** − **D**^−1*/*2^**AD**^−1*/*2^, where **D** = diag _*j*_ *A*_*ij*_ . We compute **L** = **UΛU**^⊤^ and discard the trivial eigenvector at *λ*_0_ = 0. The spectral coordinate of gene *g*_*i*_ is: **s**_*i*_ = **U**_*i*, 1:*K*+1_ ∈ ℝ^*K*^, where *K* controls the resolution of the spectral embedding. These coordinates are stored in a lookup tensor **S** ∈ ℝ^|*V*|×*K*^ indexed by token id. The eigen-decomposition is performed once before training and gene set *S* is frozen thereafter; because every training step and every inference query consults the same fixed tensor, the sign convention produced by the single decomposition is absorbed into Ω during training and no sign ambiguity arises across instances. We introduce a learnable tensor 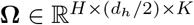, where *d*_*h*_ = *d/H*, that projects spectral coordinates to rotation angles independently per head *h* and dimension pair *n*:

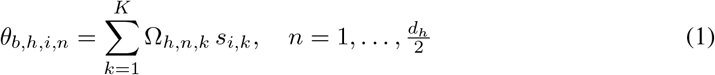

Angles enter rotary encoding [12] that rotates each adjacent pair of query & key dimensions by *θ*_*n*_:

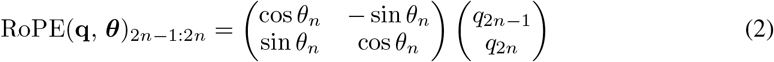

This encoding is applied within attention layers in the gene encoder to produce topology-aware gene representations. It also contributes as the spectral component of the combined rotation angle ***ψ*** applied at every backbone attention operation.

Genes with similar co-expression topology receive proximate spectral coordinates and attend to each other, while each head can specialize to a different scale of gene community structure via **Ω**, initialized to *N* (0, *K*^−1^) and learned end-to-end. Token indices are mapped to embeddings, layer-normalized, and passed through this spectral self-attention layer to produce gene representations **E** ∈ ℝ^*B*×*N* ×*d*^, which are shared between the gene identity and the perturbation embedding pathway.

### Expression and Perturbation Embeddings

Gene expression values span several orders of magnitude and contain structural zeros from dropout, which are statistically distinct from true biological absence. For expression vectors **c** ∈ ℝ^*B*×*N*^, each value is encoded as **v**_*i*_ = *f*_*ϕ*_(min(*c*_*i*_, *c*_max_)), where *f*_*ϕ*_ : ℝ → ℝ^*d*^ is a two-layer MLP with ReLU activation and layer normalization. The token representation entering the backbone is **x**_*i*_ = **v**_*i*_ + **e**_*i*_, where **e**_*i*_∈ ℝ^*d*^ is the corresponding gene embedding from the spectral encoder. Two independent value encoders 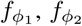, are used for the interpolated noisy state **x**_*t*_ and the conditioning control cell **c**_ctrl_ respectively. For perturbations, 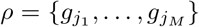 is the set of knocked-out genes. We encode the perturbation through the gene encoder with shared weights, which are then mean-pooled. Weight sharing places the perturbation representation in the same space as the gene embeddings, so the model reasons about knocked-out and measured genes through a common basis. To enable classifier-free guidance [19], **p** is replaced during training by a learned mask token **e**_mask_ ∈ ℝ^*d*^ with probability *p*_drop_. At inference, a guidance scale *ω* ≥ 1 amplifies the conditional signal: 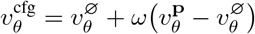

### Adaptive Repositioning (AR)

Spectral coordinates are fixed at training time from the control-state co-expression graph and cannot reflect perturbation-induced rewiring: a gene that is a hub under control conditions may become functionally isolated under a knockout. We address this through a second positional mechanism that operates at every backbone attention layer, deriving gene positions dynamically from hidden states. At layer *l*, the hidden state 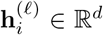 of gene *i* is projected to a scalar position per attention head via a SwiGLU gate [20]:

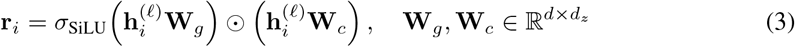

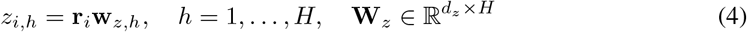

where *d*_*z*_ = max(*d/*8, 16). **W**_*z*_ is initialized from *N* (0, 0.01^2^) so the model begins training with no positional bias, allowing positions to be learned purely from data. Scalar positions are converted to rotary angles at geometric frequencies:

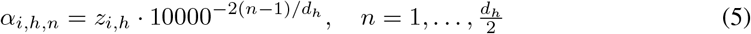

Queries and keys are rotated by *α*_*i,h*,_· before the attention dot product. By the RoPE inner product identity, the score between genes *i* and *j* in head *h* depends only on the relative position *z*_*j,h*_ − *z*_*i,h*_:

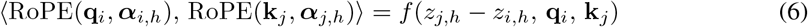

Because 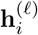 integrates gene identity, expression value, and perturbation conditioning injected at every layer via the adapter, *z*_*i,h*_ tracks each gene’s functional role under the specific perturbation at inference time without any explicit supervision on gene roles. Concretely, a hub gene knocked out in *ρ* will propagate its altered hidden state through the perturbation adapter at every layer, systematically shifting *z*_*i,h*_ away from its control-state value; Fig. 3 shows empirically that AR’s contribution over SPE grows in later training stages; we treat the hub gene z-shift as a mechanistic hypothesis consistent with this observation.

**Figure 1:**
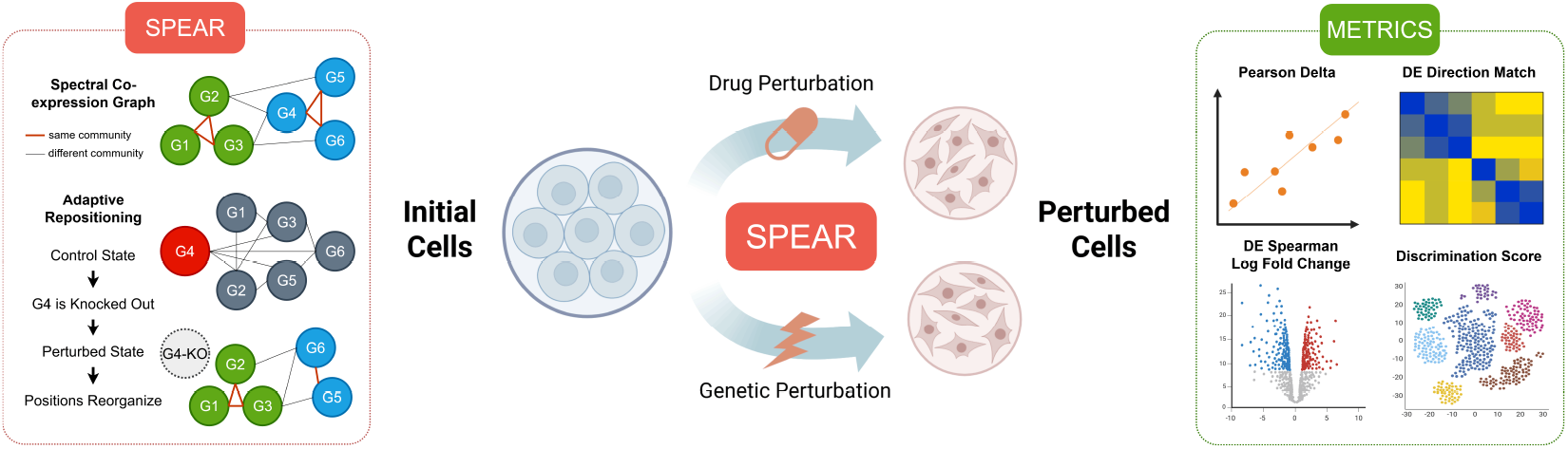
**(Left)** SPEAR’s positional encoding mechanisms illustrated by G4’s knockout collapsing hub connections and reorganizing the positional structure of remaining genes. **(Center)** SPEAR maps initial unperturbed cell populations to perturbed cell states under both drug and genetic perturbation conditions via a learned flow. **(Right)** Evaluation metrics used to assess prediction quality.

**Figure 2:**
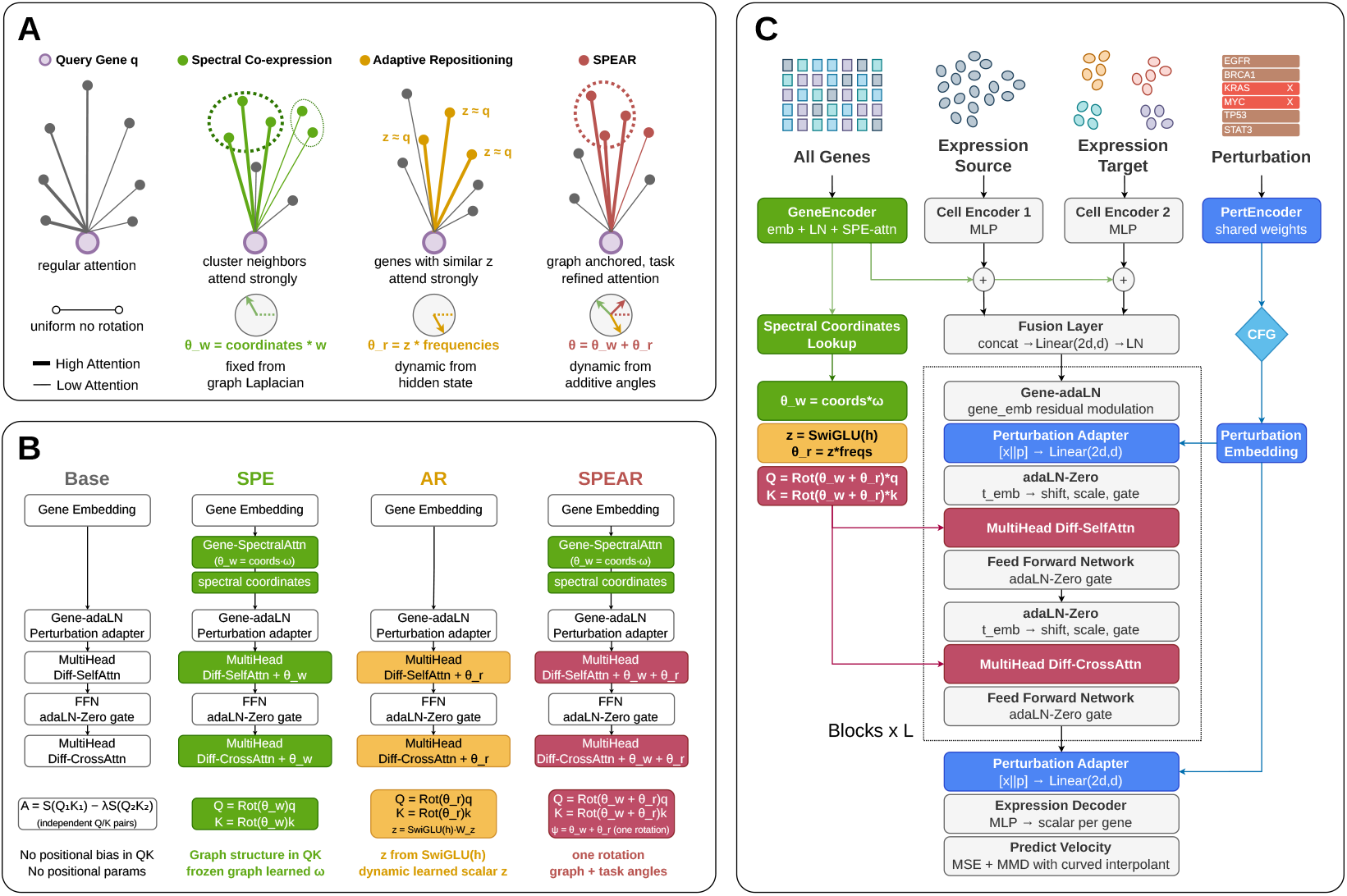
**(A)** Attention geometry under each variant. **(B)** Ablation variants showing how SPE, AR, and their combination modify attention operations. **(C)** Full model architecture pipeline of SPEAR.

**Figure 3:**
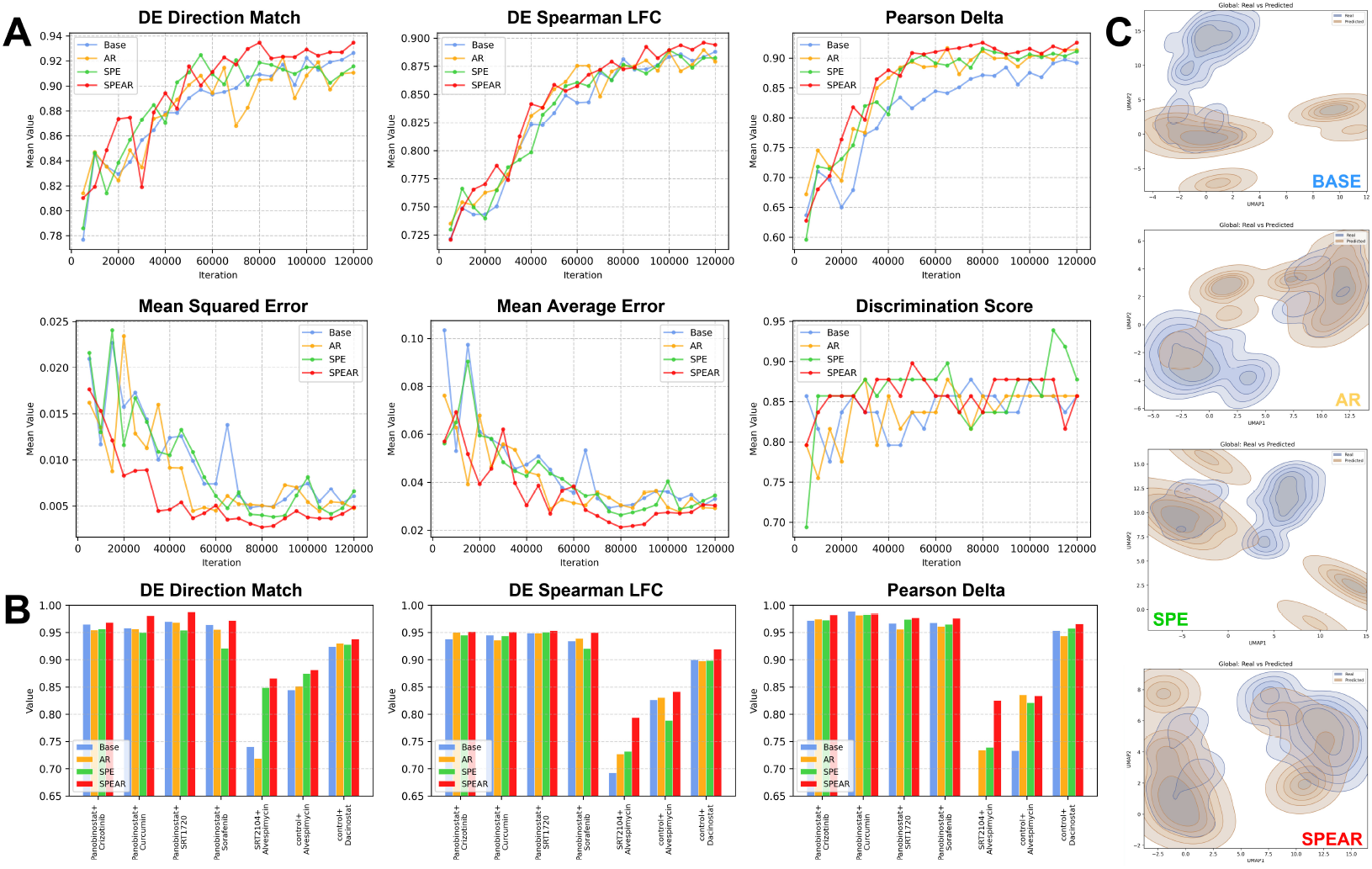
Ablation study on ComboSciPlex. **(A)** Training curves for Base, AR, SPE, and SPEAR across all evaluation metrics. **(B)** Per-drug metric breakdown on ComboSciPlex for DE Direction Match, DE Spearman-LFC, and Pearson-Δ. **(C)** Global real versus predicted UMAP density overlays for each ablation variant at iteration 120K.

### Model Architecture

The backbone processes the noisy state **Y** through *L* identical layers. Gene identity is injected via an adaptive layer norm whose shift, scale, and gate are derived from gene embeddings **E**. The perturbation embedding 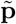 is concatenated with the hidden state and projected back to ℝ^*d*^, injecting perturbation context before any attention is computed. Each layer applies two differential attention operations [21]. First a self-attention pass that refines **Y** against itself. Second, a differential contrast pass: four independent projections produce (*Q*_*Y*_, *K*_*Y*_ ) from *Y* and (*Q*_*X*_, *K*_*X*_ ) from *X*_*c*_*trl*. Values *V* projected from *Y* only, giving **A** = softmax 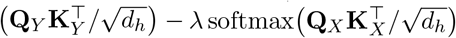, so the subtracted map cancels attention patterns shared between the noisy state and control cell, sharpening focus on their deviation. Each output is normalized by per-head RMSNorm and scaled by 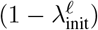. Timestep conditioning is applied via adaLN-Zero modulation derived from **t**_emb_. At every attention operation in both passes, queries and keys are rotated by the combined angle ***ψ*** = ***θ*** + ***α***, where ***θ*** is the spectral co-expression component and ***α*** is the context-adaptive component. The spectral frequency projections **Ω** are parameters at each backbone layer, allowing different layers to emphasize different scales of gene community structure.

### Training Objective

The model is conditioned on the flow time *t* ∈ [0, 1] via sinusoidal embeddings followed by a two-layer MLP. We construct training trajectories using a perturbation-conditioned interpolant that deforms the straight-line path. Given a control cell **c**_ctrl_, a noise sample **c**_0_ ∼*N* (0, **I**), and a target perturbed cell **c**_1_, a correction network ***ϕ*** bends the path:

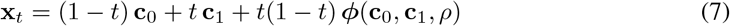

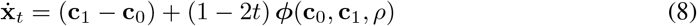

The correction vanishes at both endpoints by construction, preserving **x**_0_ = **c**_0_ and **x**_1_ = **c**_1_. ***ϕ*** is a three-layer MLP with SiLU activations taking [**c**_0_ ∥**c**_1_ ∥**e**_*ρ*_] as input, where **e**_*ρ*_ is a learned perturbation identity embedding parameterized as a lookup over training perturbations to provide perturbation-specific curvature without a full backbone forward pass. The parameters of ***ϕ*** are trained jointly with *v*_*θ*_. ***ϕ*** is discarded at inference; it shapes only the geometry of training trajectories, reducing curvature and velocity-matching variance in the same spirit as optimal transport coupling, without introducing any dependence on *c*_1_ at generation time. We sample *t* ∼*U* (0, 1), construct 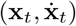 via the curved interpolant, and minimize:

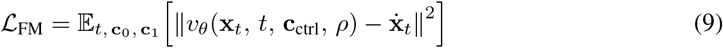

To improve distributional fidelity beyond pointwise velocity matching, we penalize distance between predicted endpoint **ĉ**_1_ = **x**_*t*_ + *v*_*θ*_ (1 − *t*) and true target **c**_1_ using an unbiased multi-scale MMD [22] estimator. Kernel bandwidths 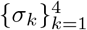 are set per batch by the median heuristic on **c**_1_:

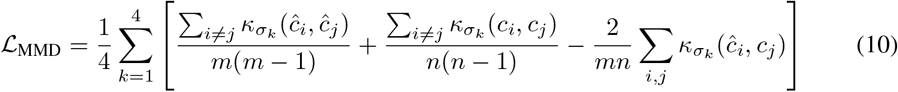

where *κ*_*σ*_(**x, y**) = exp (−∥**x** −**y** ∥^2^*/*2*σ*^2^) . The total training objective is ℒ= ℒ_FM_ + *γ ℒ*_MMD_, with *γ* a scalar hyperparameter. At inference, we integrate *v*_*θ*_ from *t* = 0 to *t* = 1 using a fixed-step RK4 solver and clamp outputs to ℝ_≥0_.

## 4 Experiments

### Baselines

We evaluate against latent-variable models, graph-based approaches, foundation models, and flow-based methods: CPA [6], GEARS [7], Geneformer [8], scGPT [9], STATE [10], CellFlow [11], LingshuCell [23], and scDFM [4]. All models take log-transformed expression as input; evaluation is restricted to the top 1,000 highly variable genes. Control returns the unperturbed profile as a lower-bound reference. CPA encodes control profiles into a latent space, adds learned perturbation embeddings, and decodes to gene space. GEARS propagates perturbation effects over a GNN-based gene interaction graph, conditioning jointly on all perturbed targets. Geneformer fine-tunes a pretrained single-cell transformer checkpoint with a lightweight perturbation prediction head. scGPT is trained from scratch as a gene-token transformer conditioned on control expression and perturbation identity. STATE conditions on basal cell state and perturbation embeddings, also trained from scratch. CellFlow learns a perturbation-conditioned flow in PCA space and decodes predictions back to gene space. LingshuCell is trained with a masking-and-prediction objective over discrete gene-expression tokens. scDFM, the closest prior to SPEAR, uses a differential transformer backbone with flow matching and a static binary co-expression attention mask. All baseline results for Combosciplex and Norman are taken from scDFM [4] and for Parse-PBMC are taken from STATE [10] as we perform the experiments on the exact same dataset splits and fixed seed.

### Datasets

The Norman dataset [24] contains genetic perturbations (CRISPR-based overexpression) in the K562 cell line, including both single and double perturbations. The ComboSciPlex dataset [25] contains drug perturbations and probes cross-modality generalization beyond genetic perturbations. The Parse-PBMC dataset [26] contains cytokine-driven transcriptomic perturbations, which profiles peripheral blood mononuclear cells. In the **additive** split, all single perturbations appearing in held-out combinations are observed during training, requiring generalization to unseen combinations of known perturbations. In the **holdout** split, selected perturbations and all combinations involving them are excluded from training, requiring extrapolation to unseen perturbation identities.

### Metrics

We evaluate across complementary aspects of prediction quality [27]. **MSE** and **MAE** measure cell-level reconstruction accuracy, with MSE emphasizing large deviations. **Discrimination Score (DS)** uses rank-based pseudobulk retrieval to assess perturbation-specific identifiability. **DE-Spearman-LFC** computes Spearman correlation between predicted and true log-fold changes restricted to differentially expressed genes selected from the ground-truth held-out profile. **Pearson-**Δ measures agreement in cell-wise gene-gene correlation structure. **Direction Match (DM)** shows if the predicted expression shift from control points in the same direction as the true perturbation vector.

### Quantitative comparison

Our consistent advantage on Spearman-LFC across all datasets is significant as it isolates recovery of biologically relevant differential expression directionality and magnitude restricted to significantly differentially expressed genes, which is the primary quantity of interest in perturbation modeling. On ComboSciPlex (Table 1), SPEAR achieves the lowest reconstruction error across all pointwise metrics and the highest biological fidelity scores. The improvement over scDFM, the closest prior work sharing the differential transformer backbone and flow matching objective, is consistent across every metric, with the largest relative gains on Spearman-LFC and Pearson-Δ, confirming that the spectral and adaptive positional encodings contribute beyond the backbone and training objective alone. On the Norman genetic perturbation benchmarks (Table 3), SPEAR achieves state-of-the-art performance across all three splits. scDFM’s marginal advantage on Pearson-Δ in the holdout splits reflects its stronger pointwise reconstruction of global expression profiles, a quantity heavily influenced by highly expressed housekeeping genes that dominate correlation statistics but are not differentially expressed under perturbation. The holdout double split, which requires extrapolation to combinations involving unobserved perturbation identities, shows the most pronounced gap over all baselines on both Spearman-LFC and DM. In the additive split, SPEAR achieves the highest Spearman-LFC across all models and splits, alongside the top DM score. On Parse-PBMC (Table 2), SPEAR leads on MSE, MAE, Spearman-LFC, and DM; gap on Pearson-Δ and DS relative to LingshuCell reflects its large-corpus pretraining that provides strong priors on population-level covariance structure.

**Table 1:** Evaluation metrics on Combosciplex.

| Model | MSE ↓ | MAE ↓ | Spear-LFC ↑ | Pears-Δ ↑ | DS ↑ | DM ↑ |
|---|---|---|---|---|---|---|
| Control | 0.0324 | 0.0698 | N.A. | N.A. | 0.5714 | 0.6167 |
| scGPT | 0.0031 | 0.0251 | -0.1261 | 0.8322 | 0.8571 | 0.8209 |
| CPA | 0.0029 | 0.0240 | 0.7906 | 0.8150 | 0.8879 | 0.8825 |
| scDFM | 0.0028 | 0.0220 | 0.8289 | 0.8933 | 0.8775 | 0.8996 |
| **SPEAR** | **0.0026** | **0.0212** | **0.8962** | **0.9259** | **0.8982** | **0.9346** |

**Table 2:** Evaluation metrics on Parse-PBMC.

| Model | MSE ↓ | MAE ↓ | Spear-LFC ↑ | Pears-Δ ↑ | DS ↑ | DM ↑ |
|---|---|---|---|---|---|---|
| scVI | 0.0045 | 0.0279 | 0.2672 | 0.0362 | 0.7263 | 0.6050 |
| scGPT | 0.0056 | 0.0737 | 0.1536 | 0.3577 | 0.7522 | 0.6825 |
| STATE | 0.0038 | 0.0261 | 0.6071 | 0.4601 | 0.8695 | 0.7214 |
| Lingshu | 0.0040 | 0.0304 | 0.5093 | **0.5925** | **0.8821** | 0.7039 |
| **SPEAR** | **0.0036** | **0.0258** | **0.6777** | 0.4614 | 0.7899 | **0.7755** |

**Table 3:**
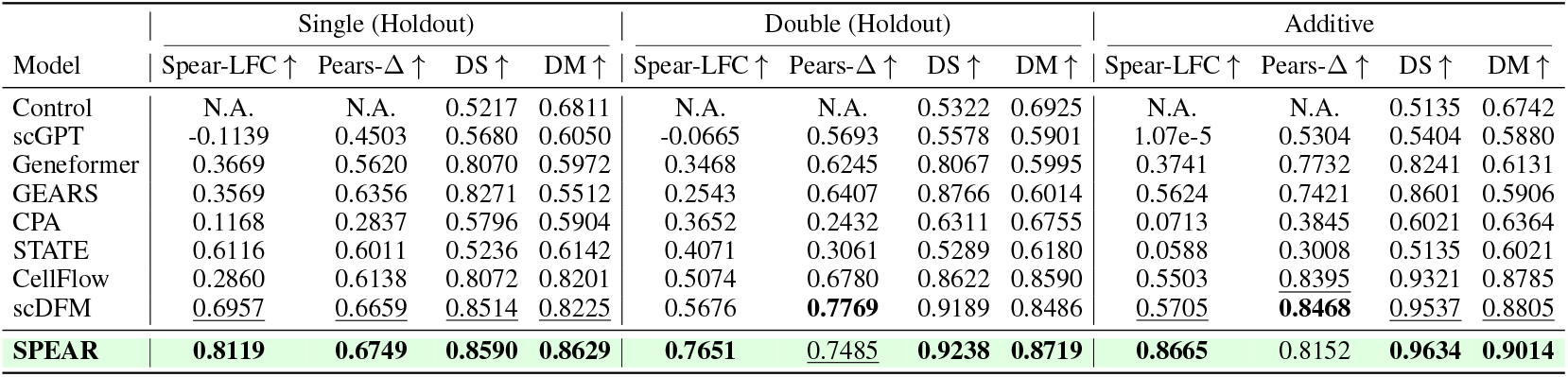
Evaluation metrics on Norman splits: holdout single, double; and additive.

| Model | Single (Holdout) |  |  |  | Double (Holdout) |  |  |  | Additive |  |  |  |
|---|---|---|---|---|---|---|---|---|---|---|---|---|
| Model | Spear-LFC ↑ | Pears-Δ ↑ | DS ↑ | DM ↑ | Spear-LFC ↑ | Pears-Δ ↑ | DS ↑ | DM ↑ | Spear-LFC ↑ | Pears-Δ ↑ | DS ↑ | DM ↑ |
| Control | N.A. | N.A. | 0.5217 | 0.6811 | N.A. | N.A. | 0.5322 | 0.6925 | N.A. | N.A. | 0.5135 | 0.6742 |
| scGPT | -0.1139 | 0.4503 | 0.5680 | 0.6050 | -0.0665 | 0.5693 | 0.5578 | 0.5901 | 1.07e-5 | 0.5304 | 0.5404 | 0.5880 |
| Geneformer | 0.3669 | 0.5620 | 0.8070 | 0.5972 | 0.3468 | 0.6245 | 0.8067 | 0.5995 | 0.3741 | 0.7732 | 0.8241 | 0.6131 |
| GEARS | 0.3569 | 0.6356 | 0.8271 | 0.5512 | 0.2543 | 0.6407 | 0.8766 | 0.6014 | 0.5624 | 0.7421 | 0.8601 | 0.5906 |
| CPA | 0.1168 | 0.2837 | 0.5796 | 0.5904 | 0.3652 | 0.2432 | 0.6311 | 0.6755 | 0.0713 | 0.3845 | 0.6021 | 0.6364 |
| STATE | 0.6116 | 0.6011 | 0.5236 | 0.6142 | 0.4071 | 0.3061 | 0.5289 | 0.6180 | 0.0588 | 0.3008 | 0.5135 | 0.6021 |
| CellFlow | 0.2860 | 0.6138 | 0.8072 | 0.8201 | 0.5074 | 0.6780 | 0.8622 | 0.8590 | 0.5503 | 0.8395 | 0.9321 | 0.8785 |
| scDFM | 0.6957 | 0.6659 | 0.8514 | 0.8225 | 0.5676 | **0.7769** | 0.9189 | 0.8486 | 0.5705 | **0.8468** | 0.9537 | 0.8805 |
| **SPEAR** | **0.8119** | **0.6749** | **0.8590** | **0.8629** | **0.7651** | 0.7485 | **0.9238** | **0.8719** | **0.8665** | 0.8152 | **0.9634** | **0.9014** |

### Qualitative cell state recovery

The global UMAP in Fig. 4A shows strong overlap between real and predicted perturbed cell distributions, with predicted contours closely tracking the multimodal structure of the true distribution. At the perturbation level, cases MAP2K3+MAP2K6 and ETS2+MAP7D1 show tight alignment of predicted and real density modes. Challenging perturbations FOXA3+FOXL1 and FOXA3+FOXL2 reveal partial misalignment in secondary density modes, consistent with lower per-perturbation scores visible in the metric distributions of Fig. 4C. Expression delta heatmaps in Fig. 4B confirm that predicted fold-change patterns correlate qualitatively with true patterns across perturbations both in presence and absence of outlier perturbations, with the without-outliers panels showing tighter correspondence in the moderate fold-change regime. Beyond this global visual alignment, SPEAR’s perturbation-responsive geometry yields actionable biological fidelity: predicted TF-perturbation responses concentrate on the factor’s TRRUST targets relative to a degree-matched null (Fig. 6B), and predicted expression shifts recover significantly more ground-truth pathway enrichments across perturbations (Enrichr p=0.004, GSEA p=0.008; Fig. 6C).

**Figure 4:**
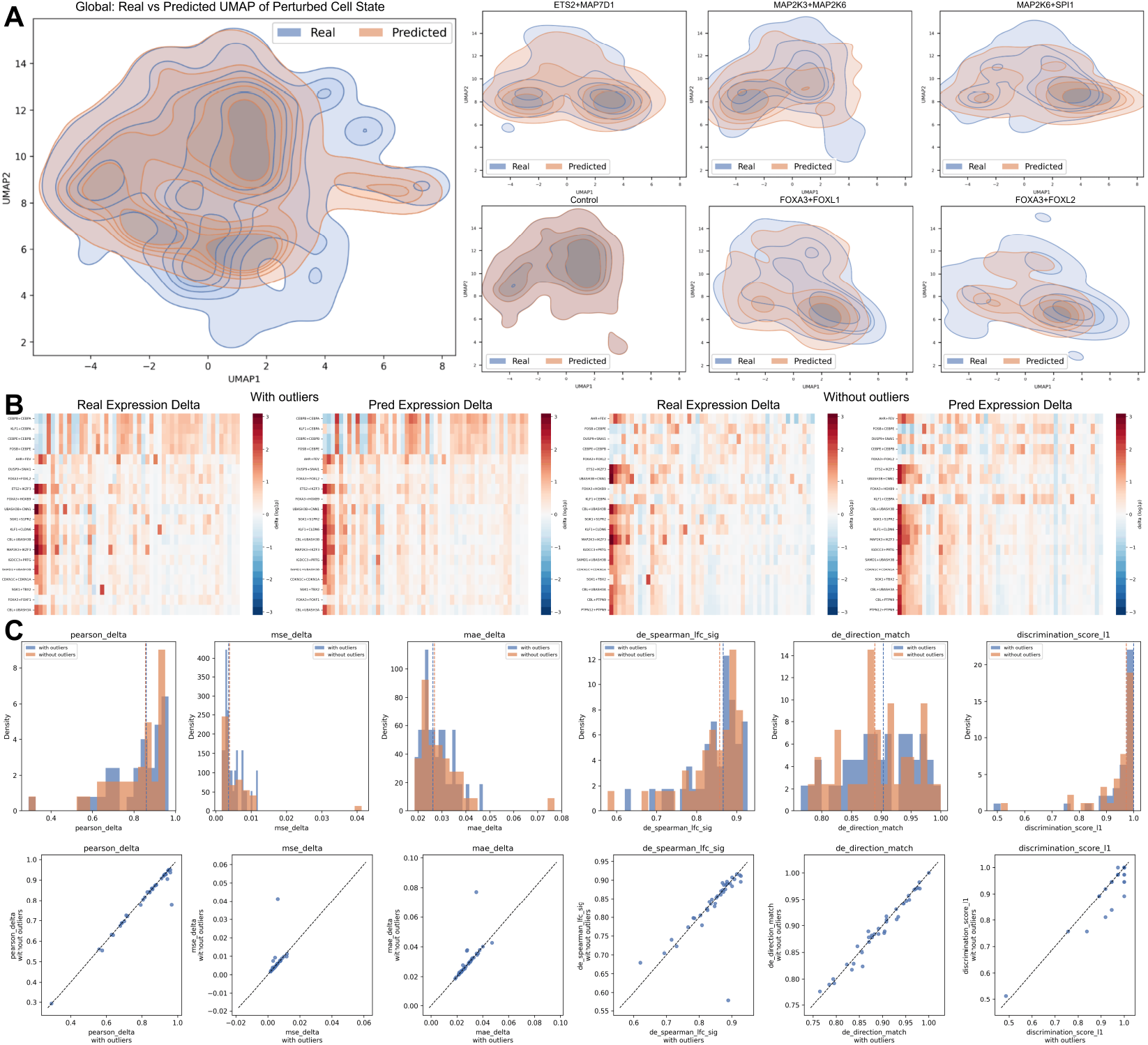
Prediction quality on the Norman dataset. **(A)** Global and per-perturbation UMAP density overlays of real versus predicted perturbed cell states. **(B)** Expression delta heatmaps across perturbations with and without outliers, showing predicted versus true log fold-change patterns. **(C)** Per-perturbation metric distributions and with-versus-without-outlier scatter plots across Pearson-Δ, MSE, MAE, DE Spearman-LFC, DM, and DS.

### Ablation study

To isolate the contribution of each positional encoding mechanism, we evaluate four variants: Base (transformer with no positional encoding beyond token embeddings), AR (context-adaptive repositioning only), SPE (spectral positional encoding only), and SPEAR (both). On the ComboSciPlex data, the training curves in Fig. 3A show convergence behavior, and SPE produces the largest early improvement over Base across DM, Spearman-LFC, and Pearson-Δ, consistent with the spectral prior providing a geometric inductive bias that accelerates convergence. AR yields additional gains over SPE in the later training phase across all three metrics, and SPEAR consistently achieves the highest values at convergence. On reconstruction error, SPEAR reaches lower MSE and MAE than all ablations throughout training. The per-drug breakdown on ComboSciPlex in Fig. 3B shows that SPEAR’s advantage over SPE and AR is consistent across individual drug combinations rather than driven by a subset of conditions, confirming that the two mechanisms are complementary. The qualitative UMAPs in Fig. 3C illustrate this directly: Base produces poorly aligned predicted distributions, AR partially recovers the global structure but misses fine-grained cluster geometry, SPE aligns broad density modes, and SPEAR achieves the tightest overall overlap with the real distribution. Fig. 5A evaluates CFG dropout rates of 0%, 5%, and 10% during training on Norman dataset. Zero dropout achieves the lowest MSE and MAE throughout training, as the model is never required to generate unconditional predictions and full capacity is directed toward conditional reconstruction. However, 5% and 10% dropout produce higher DS, DM and Spearman-LFC at convergence, indicating that CFG training improves perturbation-specific identifiability of predictions even at a cost to pointwise reconstruction error. A model can minimize MSE/MAE by regressing to the mean while producing zero differential signal; DS, DM, Spearman-LFC, Pearson-Δ measure only the condition-specific deviation from control, which is the actual prediction target. The 5% and 10% curves are indistinguishable on biological fidelity metrics, suggesting diminishing returns beyond 5% dropout and that a small CFG dropout rate is sufficient to obtain the discriminability benefit.

**Figure 5:**
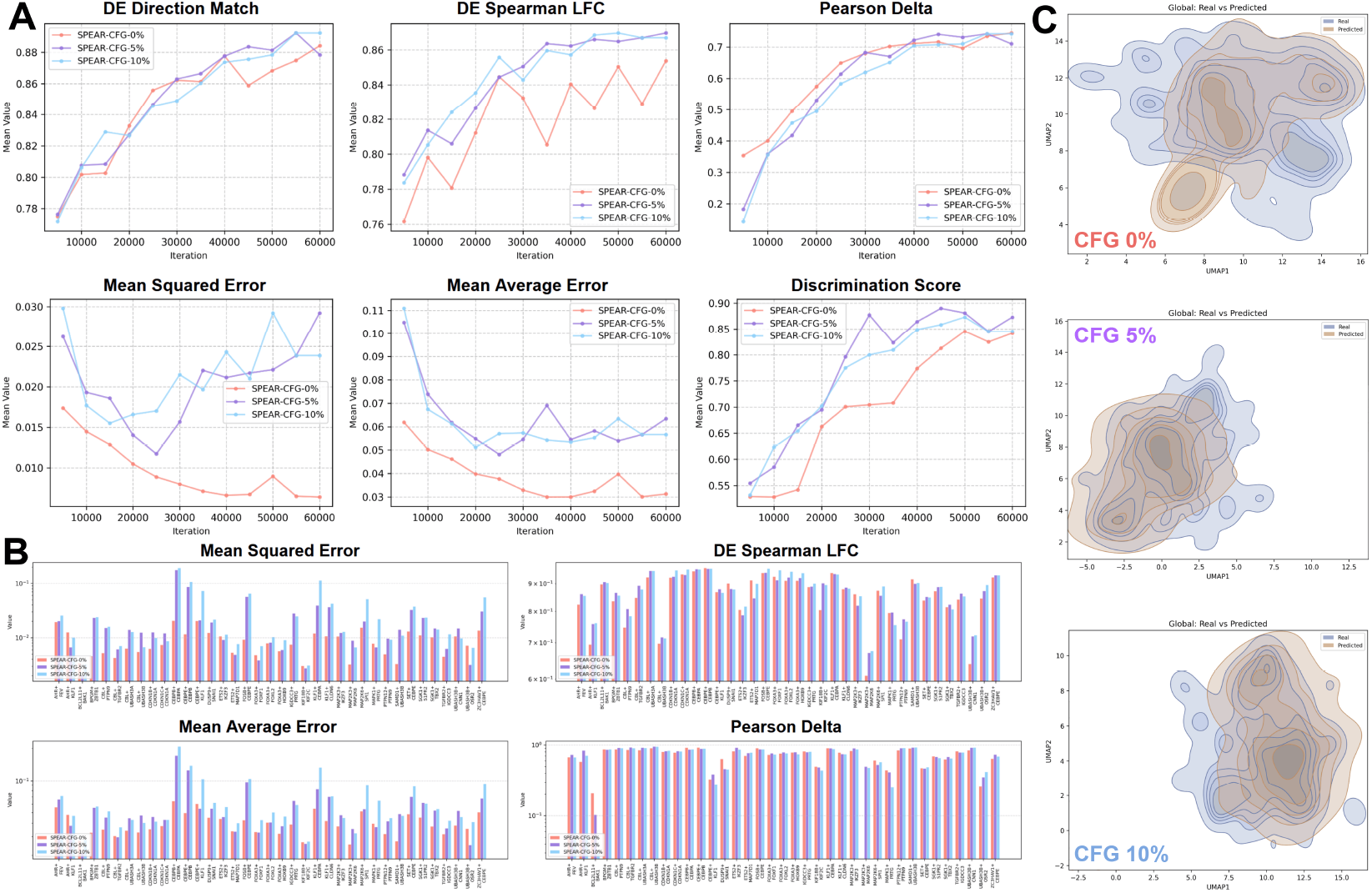
Effect of CFG dropout rate on training dynamics. **(A)** Training curves for SPEAR trained with CFG dropout rates of 0%, 5%, and 10%. **(B)** Per-drug metric breakdown on Norman for DM, DE Spearman-LFC, Pearson-Δ, MSE, MAE, and DS. **(C)** Global real versus predicted UMAP density overlays for each CFG variant at iteration 60K. We chose 5% CFG setting for final runs.

**Figure 6:**
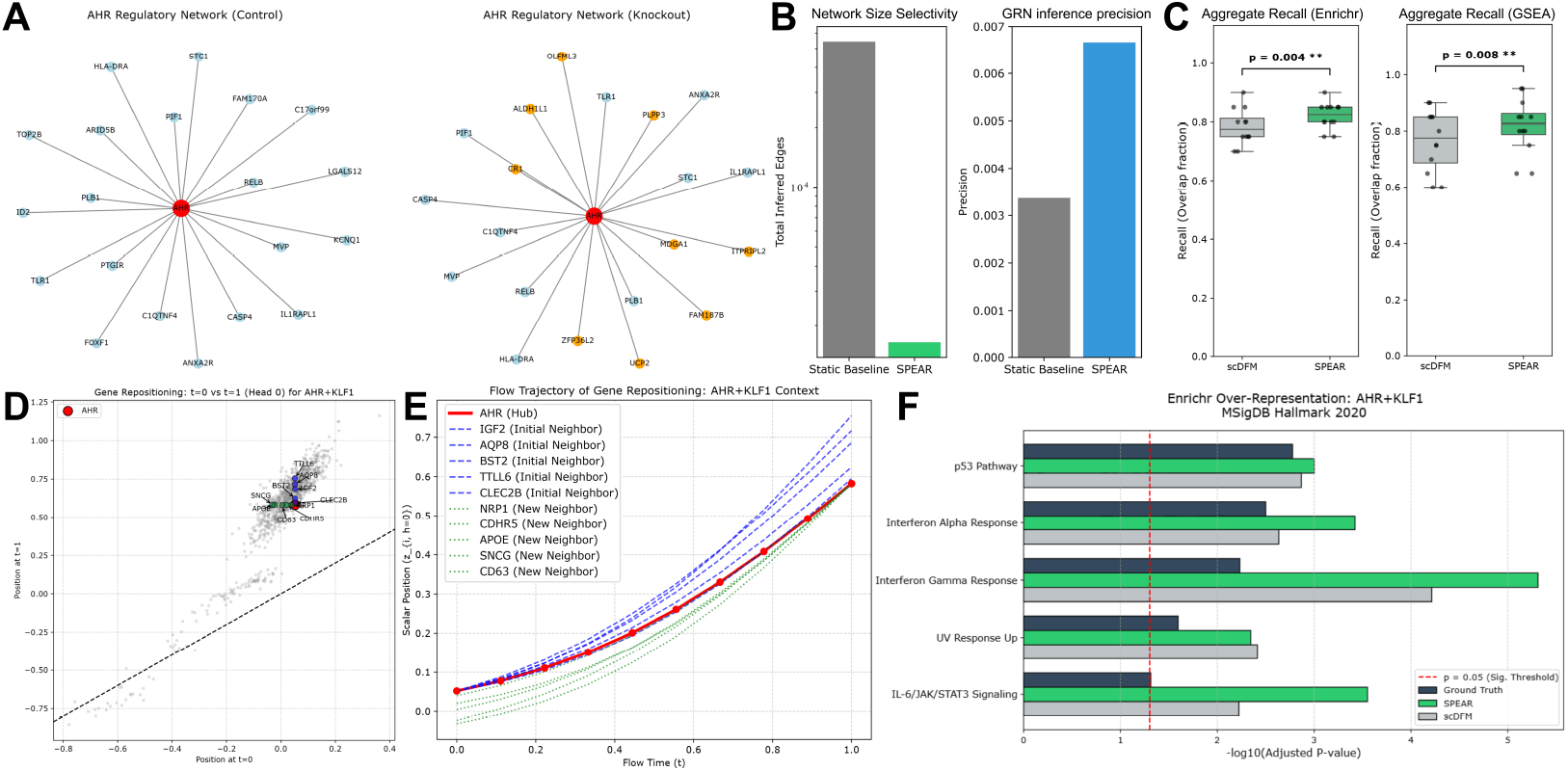
Biological interpretability of SPEAR. **(A)** AHR positional-proximity network under control vs. AHR+KLF1 knockout; retained (light blue) and rewired (orange) edges. **(B)** Sparsity (left) and TRRUST precision (right) of inferred GRNs, SPEAR vs. the static scDFM co-expression baseline. **(C)** Top-20 pathway recall across all perturbations for Enrichr (left) and GSEA (right); paired Wilcoxon tests. **(D)** Scalar position *z*_*i,h*=0_ at *t* = 0 vs. *t* = 1 under AHR+KLF1. **(E)** Flow-time position trajectories of AHR and its dynamic neighbors. **(F)** Enrichr enrichment for AHR+KLF1 on MSigDB Hallmark: SPEAR, ground truth, and scDFM.

### Biological Analysis

To test whether Adaptive Repositioning (AR) encodes meaningful gene relationships, we extracted scalar positions *z*_*i,h*_ from the first cross-attention query module at 175K iterations (Norman additive split). Since ∥*z*_*i*_− *z*_*j*_∥ modulates attention scores via the RoPE identity, we use it as an unsupervised edge score. Fig. 6A compares AHR’s top-20 positional neighbors under control vs. AHR+KLF1 knockout, evaluating the same trained model on each cell subset.

Without edge supervision, some connections are retained (light blue) and new ones form (orange), so perturbation-conditioned hidden states reconfigure neighborhoods. Against TRRUST (1000-HVG universe), the static baseline uses all edges incident to known TFs in the scDFM co-expression mask, while SPEAR uses the top-30 AR neighbors per TF under control. With orders of magnitude fewer edges, SPEAR achieves roughly double the precision (Fig. 6B), as the dense mask conflates indirect co-regulation and technical covariation with direct interactions. Thus AR distances, though not trained for GRN inference, reflect biologically coherent proximity. SPEAR outperforms scDFM in recalling the top-20 altered pathways across all perturbations (*p* = 0.004 Enrichr, *p* = 0.008 GSEA; Fig. 6C). For AHR+KLF1 on MSigDB Hallmark 2020 (Fig. 6F), SPEAR recovers higher− log_10_ adjusted *p* for core responses such as p53 and Interferon Gamma, which scDFM underestimates. Plotting *z*_*i,h*=0_ at *t* = 0 vs. *t* = 1 under AHR+KLF1 (Fig. 6D), most genes lie on the diagonal while AHR and its neighborhood deviate consistently, indicating perturbation-driven shifts rather than noise. Across 10 uniformly spaced flow timesteps (Fig. 6E), trajectories evolve smoothly and non-linearly: initial neighbors track the hub early then diverge, and new neighbors converge toward it late. Positional proximity is therefore dynamically constructed and perturbation-conditioned.

## 5 Discussion

SPEAR demonstrates that positional encoding is a key bottleneck for perturbation prediction in gene transformers. The encoding contribution is isolated by controls within our pipeline rather than by comparison to scDFM: permuted and freely learned coordinates of the same dimensionality do not reproduce SPEAR’s DE Spearman-LFC gain (Table 5). The largest gains on Spearman-LFC suggest that spectral and adaptive encodings improve resolution of gene-specific responses, while MSE/MAE gains reflect global expression magnitude. Ablations show an asymmetry: SPE is a structural inductive bias that reduces the optimizer’s search space, while AR contributes later in training by refining perturbation-specific geometry the fixed spectral prior cannot represent. The co-expression graph encodes stable control-state relationships and adaptive positions add perturbation-conditioned corrections, so the two mechanisms operate at different levels of the positional hierarchy and are complementary by construction. Norman holdout splits demand generalization to unseen perturbation combinations. scDFM’s marginal Pearson-Δ advantage reflects reconstruction of background expression rather than the perturbation-specific program, making Spearman-LFC, restricted to significantly differentially expressed genes, the more biologically meaningful metric. SPEAR’s advantages on Spearman-LFC and all other metrics suggest adaptive positional encoding is particularly valuable under distribution shift: since scalar positions derive from perturbation-conditioned hidden states, the model can reconfigure gene influence for novel knockouts without relying on a fixed graph. Consistent state-of-the-art performance across three perturbation modalities, split regimes, and evaluation criteria establishes biologically grounded positional encoding as a principled and necessary component of transcriptomic perturbation modeling. **Limitations**. Spectral coordinates are precomputed from the training co-expression graph and cannot accommodate genes outside the training vocabulary. Adaptive positions are computed in a single forward pass and cannot use feedback from downstream layers when computing upstream rotations. Whether SPEAR retains its advantage under noisier co-expression estimates at full transcriptome scale remains open, though the demonstrated gains motivate more robust graph estimation. **Future Work**. An obvious extension would be multi-modal perturbation modelling [28] and generating perturbed cellular phenotypes in tissues [29, 30].

**Table 4:** Overview of key single-cell perturbation and reference datasets. The **ComboSciPlex** dataset provides a large-scale combinatorial drug screen utilizing A549 cells to capture complex, multi-drug transcriptional responses. The **Norman** dataset serves as a standard foundational benchmark for modeling CRISPR-mediated genetic activation in K562 cells. The **Parse PBMC** datasets represent massive-scale transcriptomic profiling using split-pool barcoding, encompassing up to 10 million peripheral blood mononuclear cells.

| **Dataset** | **Perturbation Type** | **Cell Line / Tissue** | **Total Cells** | **Conditions** |
|---|---|---|---|---|
| **ComboSciPlex** | Chemical Small Molecules | A549 (Lung Cancer) | 63,378 | 32 |
| **Norman** | Genetic CRISPRa | K562 (Leukemia) | > 100, 000 | 237 |
| **Parse PBMC** | Cytokines | Human PBMCs | ∼9.7 – 10M | 90+ |

**Table 5:** Multi-seed ablation and positional controls on ComboSciPlex (5 seeds, mean ±95% CI). Permuted and Learned replace the spectral coordinate table in SPEAR; MSE not run for controls.

| Model | MSE (×10<sup>−3</sup>) ↓ | MAE (×10<sup>−2</sup>) ↓ | Spear-LFC ↑ | Pears-Δ ↑ | DS ↑ | DM ↑ |
|---|---|---|---|---|---|---|
| Base | 2.79±0.16 | 2.08±0.08 | 0.8478±0.0111 | 0.9012±0.0166 | 0.8776±0.0179 | 0.9080±0.0074 |
| SPE | 2.64±0.42 | 2.00±0.12 | 0.8753±0.0161 | 0.9034±0.0619 | 0.8816±0.0278 | 0.9159±0.0223 |
| AR | 2.30±0.36 | 1.96±0.06 | 0.8841±0.0132 | 0.9164±0.0081 | 0.8694±0.0227 | 0.9260±0.0059 |
| Permuted | – | 1.96±0.10 | 0.8416±0.0127 | 0.9225±0.0282 | **0.8880**±0.0194 | 0.9313±0.0109 |
| Learned | – | **1.92**±0.10 | 0.8586±0.0155 | 0.9218±0.0203 | 0.8780±0.0185 | 0.9315±0.0116 |
| **SPEAR** | **2.09**±0.29 | 1.95±0.10 | **0.8984**±0.0151 | **0.9420**±0.0313 | 0.8878±0.0200 | **0.9372**±0.0202 |

## A SPEAR Training Dynamics

To confirm that both mechanisms learn structured representations rather than drifting arbitrarily, we track **Ω** (SPE) and {**W**_*g*_, **W**_*c*_, **W**_*z*_} (AR) across all checkpoints on the Norman run, computing Frobenius drift from initialization, RIP graph alignment statistics (delta mean RIP and Mann-Whitney U p-value between mask-connected and unconnected gene pairs), and head diversity metrics.

### SPE learns graph-structured frequency projections

Fig. 8A shows **Ω** drifting monotonically from its near-identity initialization, saturating after ≈75–100K iterations in line with metric convergence in Fig. 3. Cross-attention layers drift further than self-attention layers at the same depth, and deeper blocks drift further than shallower ones, reflecting the larger positional discrepancy cross-attention must resolve when aligning query states at noise level *t* against control-state keys. The encoder **Ω** (purple) barely moves, confirming it acts as a fixed spectral lookup with negligible gradient. Fig. 8D shows that each head’s **Ω**_*h*_ matrix evolves from near-diagonal at iteration 0 to distinct off-diagonal structure by 20K–30K iterations, with different heads concentrating weight on low-versus high-frequency eigenvector components, realizing the intended per-head specialization to different scales of gene community structure. RIP alignment statistics confirm that mask-connected gene pairs receive statistically significantly higher RIP scores than unconnected pairs well before convergence (positive delta mean RIP, MWU *p <* 0.05), verifying that **Ω** encodes graph proximity into rotation angles rather than learning an arbitrary projection. The RIP graph panels (Fig. 8C) show this directly: the top-*k* RIP graph transitions from diffuse at iteration 0 to a stable high-RIP core by 40K–70K, mirroring the saturation of the drift curve.

**Figure 7:**
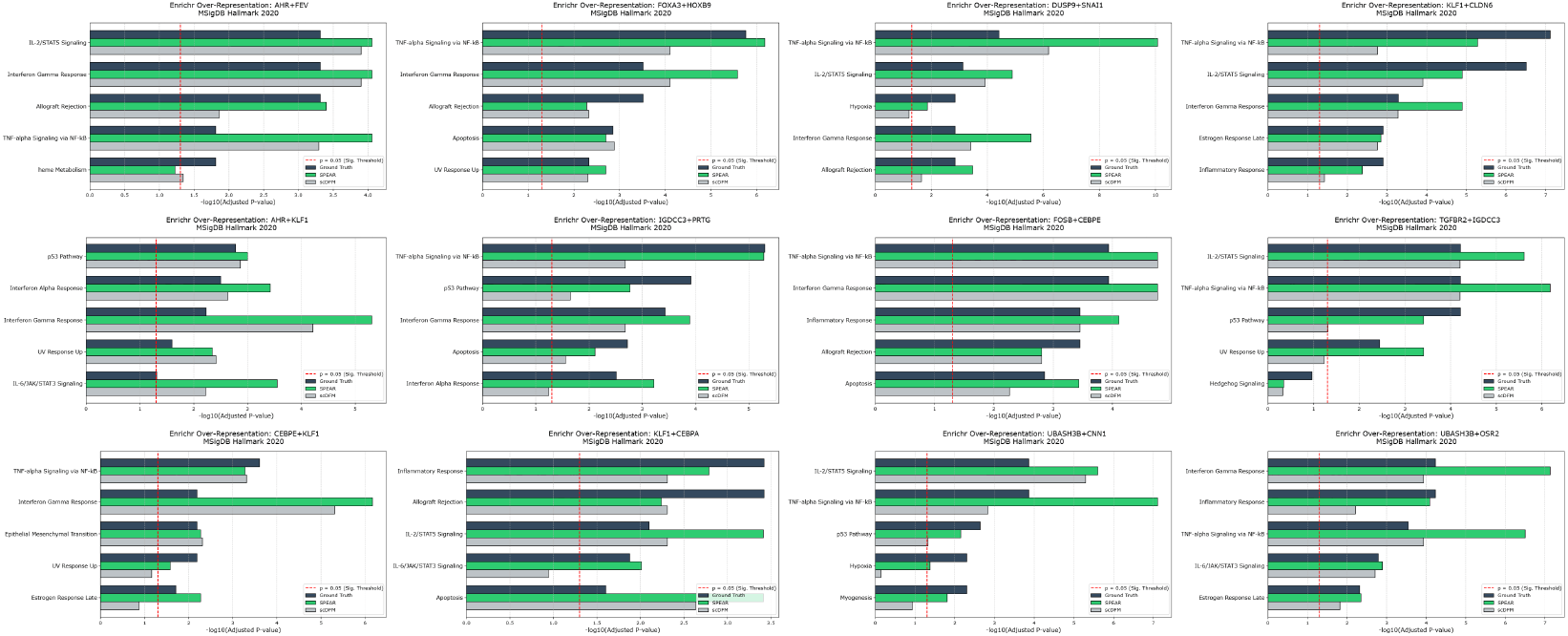
Enrichr enrichment for all Norman validation perturbations on MSigDB Hallmark.

**Figure 8:**
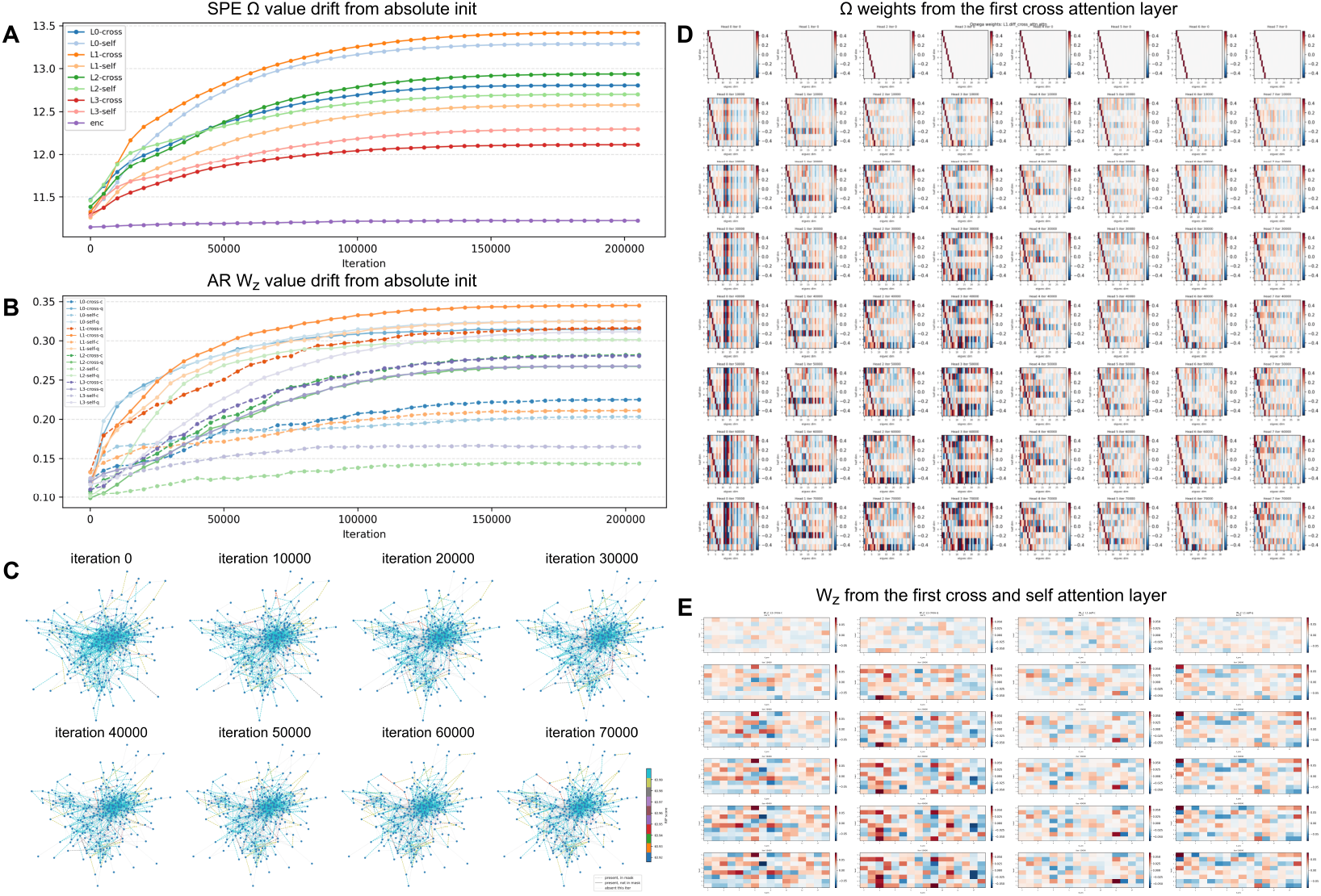
Parameter evolution of SPE and AR across training checkpoints on Norman. **(A)** Frobenius drift of **Ω** from its identity-like initialization (*N* (0, *K*^−1^)) across all backbone layers. Cross-attention layers consistently drift further than self-attention layers at the same depth, and deeper layers drift more than shallower ones. The encoder **Ω** (purple) remains nearly stationary throughout training. **(B)** Frobenius drift of **W**_*z*_ from its near-zero initialization (*N* (0, 0.01^2^)) for all adaptive repositioning groups across backbone layers. Solid lines denote query and dashed lines context; query-side weights consistently drift further than context-side weights within the same block, indicating asymmetric plasticity. Cross-attention groups again show higher drift than self-attention groups. **(C)** Spectral co-expression graph evolution showing top-*k* RIP gene pairs at milestone iterations. At initialization the graph is nearly uniform; by 10K–20K structured clusters emerge; by 40K–70K the graph stabilizes with a sparser periphery. **(D) Ω** weight heatmaps (half× eigvec_dim) per attention head in the first cross-attention layer at milestone iterations. Rows represent the half-dimension index, columns the eigenvector dimension. Near-diagonal structure at iteration 0 reflects the identity-like initialization; by 30K strong off-diagonal structure emerges with each head developing a distinct mixing pattern over eigenvector components, demonstrating head specialization to different scales of co-expression community structure. **(E) W**_*z*_ heatmaps (nhead× *d*_*z*_) for the first cross-attention and self-attention layers at milestone iterations. Cross-attention **W**_*z*_ develops more differentiated row patterns across heads relative to self-attention, consistent with cross-attention bearing greater positional alignment responsibility between the noisy state & control cell.

**Figure 9:**
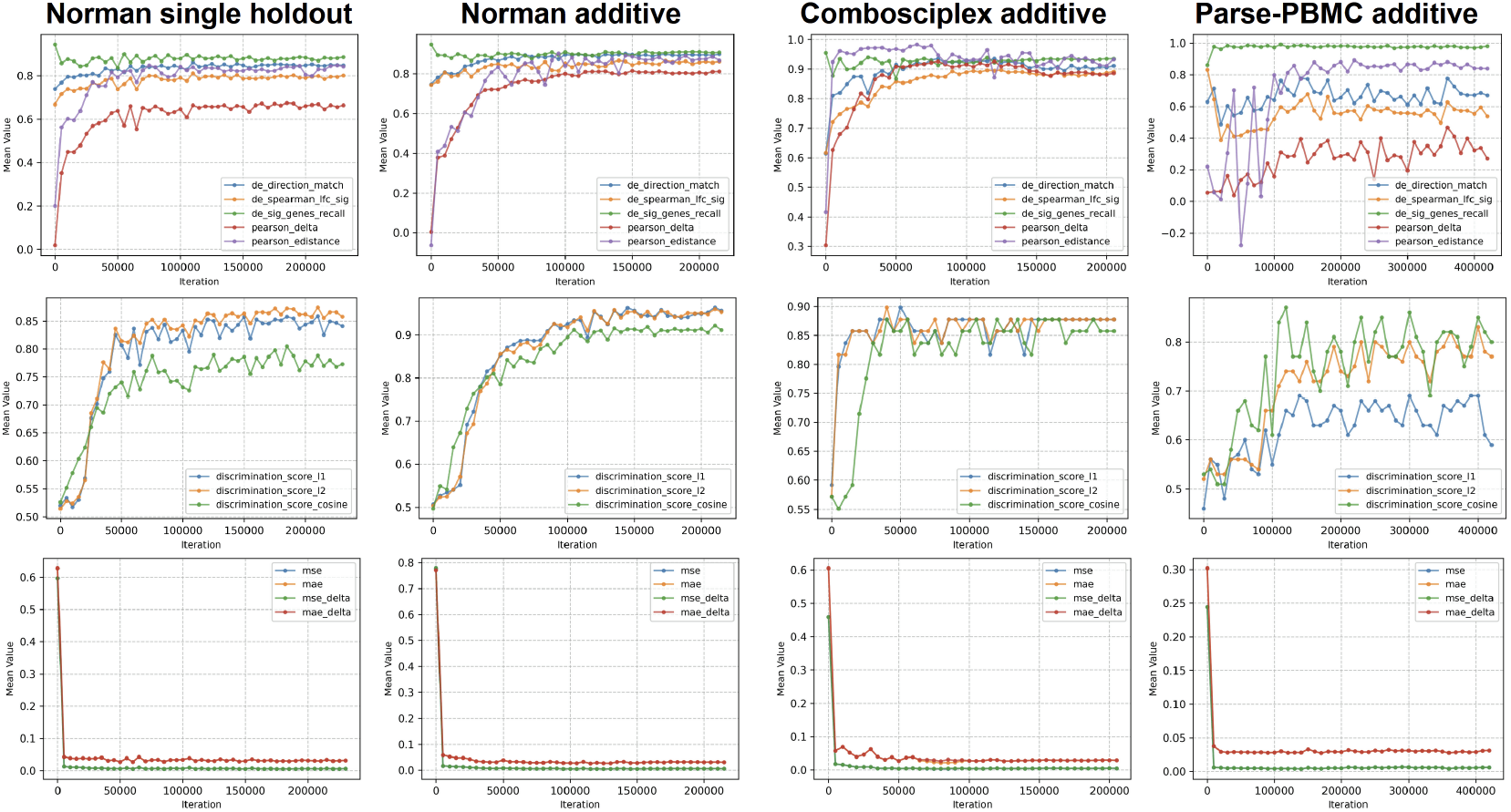
Validation metric curves on best fold for all datasets across training iterations.

### AR exhibits asymmetric query-context plasticity

Fig. 8B shows **W**_*z*_ drifts monotonically from its near-zero initialization, but query-side weights (solid lines) consistently drift further than context-side weights (dashed lines) across all blocks and attention types. This asymmetry follows directly from the training setup: query positions are computed on the interpolated noisy state **x**_*t*_, which varies with *t* throughout each training trajectory and therefore drives larger positional variance; context positions are computed on the fixed control cell **c**_ctrl_ and require less parameter movement. Fig. 8E confirms that cross-attention **W**_*z*_ develops more differentiated head row patterns than self-attention **W**_*z*_, consistent with cross-attention bearing greater positional alignment responsibility.

### Two-phase learning and complementarity

**Ω** saturates earlier than **W**_*z*_, and SPE dominates the early training gain in Fig. 3A while AR contributes most in the later phase. This ordering reflects two levels of the positional hierarchy: the co-expression Laplacian prior is rapidly encoded by **Ω** in early training, reducing the optimizer’s effective search space; once saturated, **W**_*z*_ continues learning perturbation-conditioned corrections that the fixed spectral prior cannot represent. This two-phase dynamic provides mechanistic support for the ablation finding that SPE and AR are complementary rather than redundant, and that their combination is strictly better than either alone at convergence.

### Multi-seed ablation and positional controls

All variants are retrained on ComboSciPlex over five seeds on the fixed split and read at 200K iterations (Table 5, mean ±95% CI). SPEAR is best on all six metrics, and the ordering Base *<* SPE *<* AR *<* SPEAR holds on five of six. Significance is assessed across the seven held-out perturbations after averaging seeds within each: SPEAR exceeds Base, SPE, and AR on DE Spearman-LFC in 7/7 perturbations (Δ = +0.0305, +0.0231, +0.0143), each at the minimum attainable exact one-sided Wilcoxon signed-rank *p* = 2^−7^ = 0.0078. The five secondary metrics are also unanimous (Holm-adjusted *p* = 0.0391, the minimum this design allows); since every raw *p* sits at the design floor, effect size and unanimity carry the evidence. To separate co-expression geometry from generic gene-specific rotation, we replace **S** in SPEAR with a random row permutation across genes (Permuted) or free learnable coordinates of dimension *K* (Learned), holding all other components, parameter count, and training fixed. On DE Spearman-LFC both fall 0.040–0.057 below SPEAR (2.6–3.8 times its seed interval), with Permuted below Base. On MAE, DS, and DM both controls recover most of the gain over Base, and on Pearson-Δ about half; Permuted matches SPEAR on DS. These metrics therefore do not separate co-expression geometry from arbitrary gene-specific rotation, and the geometry-specific gain is confined to DE Spearman-LFC.

### Seed variability against scDFM

Over five seeds, SPEAR’s DE Spearman-LFC margin over scDFM exceeds the larger of the two 95% seed intervals by at least 4.5× on every dataset and split (Table 6). Sweeping *K* ∈ {8, 16, 32, 64, 128} (one run per point) changes ComboSciPlex DE Spearman-LFC by 0.0156 in total (0.898–0.914), comparable to one configuration’s seed interval (± 0.0151); no oversmoothing regime appears up to *K* = 128.

**Table 6:** DE Spear-LFC over 5 seeds (mean *±* 95% CI)

| Dataset / split | scDFM | SPEAR | Margin |
|---|---|---|---|
| ComboSciPlex | 0.8296±0.0129 | 0.8984±0.0151 | +0.0688 |
| Norman single | 0.6985±0.0113 | 0.8119±0.0176 | +0.1134 |
| Norman double | 0.5677±0.0165 | 0.7651±0.0101 | +0.1974 |
| Norman additive | 0.5769±0.0185 | 0.8665±0.0156 | +0.2896 |

**Table 7:** Hyperparameters used for all reported experiments.

| Hyperparameter | Norman | ComboSciPlex / Parse-PBMC |
|---|---|---|
| *d*<sub>model</sub> | 128 | 128 |
| Attention heads *H* | 8 | 8 |
| Backbone layers *L* | 8 | 8 |
| MLP ratio | 4 | 4 |
| Eigenvector dimension *K* | 32 | 32 |
| Top-*k* neighbors | 30 | 30 |
| Highly variable genes | 5000 | 5000 |
| Genes per forward pass | 1000 | 1000 |
| Batch size | 48 | 48 |
| Optimizer | Adam | Adam |
| Learning rate | 5 × 10<sup>−5</sup> | 5 × 10<sup>−5</sup> |
| LR schedule | Cosine annealing | Cosine annealing |
| *η*<sub>min</sub> | 1 × 10<sup>−6</sup> | 1 × 10<sup>−6</sup> |
| Training steps | 200K | 200K |
| MMD loss weight *γ* | 0.5 | 0.5 |
| MMD kernel scales | {0.5, 1.0, 2.0, 4.0} | {0.5, 1.0, 2.0, 4.0} |
| Noise type | Gaussian | Gaussian |
| CFG dropout *p*<sub>drop</sub> | 0.05 | 0.05 |
| CFG guidance scale *ω* | 1.5 | 1.5 |
| ODE solver | RK4 | RK4 |
| ODE steps (inference) | 20 | 20 |
| Split method | holdout / additive | combinations |

The DE gene set for Spearman-LFC is selected from the ground-truth held-out profile (Wilcoxon against control) and shared across models, so the metric measures fold-change recovery conditional on a known DE set. With the set selected from predictions instead, DE-gene recall is 0.883± 0.0137 vs 0.868 ±0.0194 (Base) on Norman and 0.938 ±0.0171 vs 0.927 ±0.0120 on ComboSciPlex, within seed intervals: SPEAR is not worse at identifying responding genes, but no advantage is claimed. For single-TF perturbations, predicted |Δ| on the TF’s TRRUST targets relative to a degree-matched random null has mean enrichment 1.90 (median 1.69) vs 1.03 for the null (ratio 1.85), above the null in 7 of 9 qualifying TFs; the predicted sign matches the annotated direction for 52% of targets vs 43% under the null (Fig. 6B). AR proximity neighbors are reported as a property of the learned geometry, not as inferred regulatory rewiring.

## B SPEAR Mathematical Formulation

### B.1 Spectral Co-expression Encoding

#### Graph construction

Let **X** ∈ℝ^*C*×*G*^ be the log-normalized training expression matrix over *C* cells and *G* highly variable genes. Column-normalize to zero mean and unit *l*_2_ norm to obtain 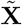, then form the symmetric co-expression adjacency:

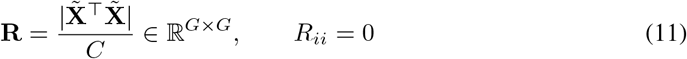

where the elementwise absolute value treats co- and anti-correlated gene pairs symmetrically. Sparsify each row to its top-*k* entries and symmetrize:

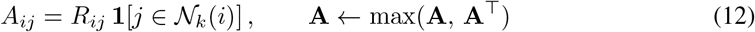

#### Laplacian eigenvectors

Form the symmetric normalized Laplacian **L** = **I** − **D**^−1*/*2^**AD**^−1*/*2^, where **D** = diag(Σ_*j*_ *A*_*ij*_). Eigendecompose **L** = **UΛU**^⊤^ via `numpy.linalg.eigh`. Discard the trivial eigenvector at *λ*_0_ = 0. The spectral coordinate of gene *g*_*i*_ is:

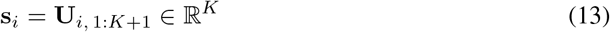

stored in a frozen lookup tensor **S** ∈ ℝ^|*V*|×*K*^ indexed by vocabulary token id. This computation is a one-time preprocessing step; the fixed sign convention from the single decomposition is absorbed into **Ω** during training.

#### Learnable frequency projection

A learnable tensor 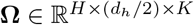 projects spectral coordinates to rotation angles independently per head *h* and dimension pair *n*:

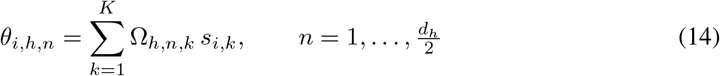

implemented as `torch.einsum(‘bge,hfe->bhgf’, coords, omega)`. **Ω** is initialized with a near-diagonal structure placing unit weight on eigenvector component *f* mod *K* for dimension pair *f*, then learned end-to-end. Each head thereby specializes to a different scale of gene community structure.

### Rotary application

Rotation angles are applied to queries and keys via the standard RoPE rotation on adjacent dimension pairs (2*n* − 1, 2*n*):

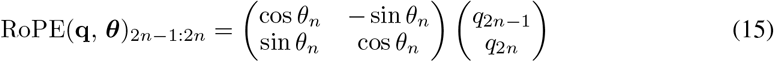

By the RoPE inner product identity, the attention score between genes *i* and *j* in head *h* depends only on the pairwise angle difference *θ*_*j,h,n*_ −*θ*_*i,h,n*_, which is a linear function of the difference in spectral coordinates **s**_*j*_− **s**_*i*_. Genes with similar co-expression topology have small spectral distance and thus high attention affinity.

#### Algorithm 1

Spectral Coexpression encoding (per attention operation)

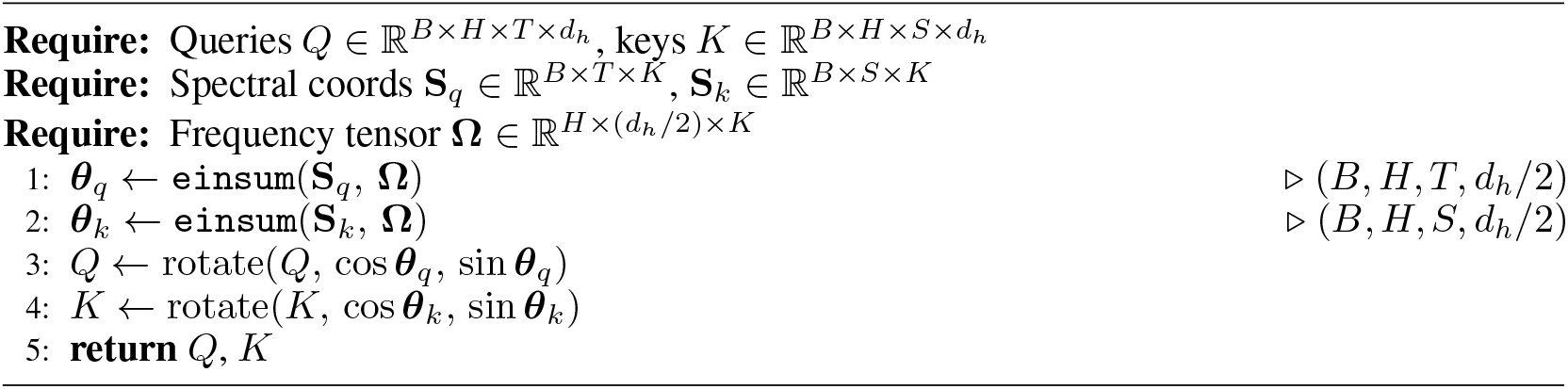

### B.2 Adaptive Repositioning

#### Scalar position derivation

At backbone layer *l*, the hidden state 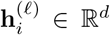 of gene *i* which integrates gene identity, expression value, and perturbation conditioning is projected to a scalar position per attention head via a SwiGLU gate:

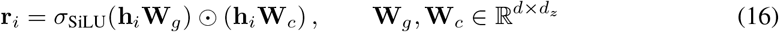

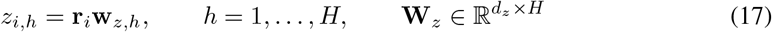

where *d*_*z*_ = max(*d/*8, 16). **W**_*z*_ is initialized from *N* (0, 0.01^2^) so the model begins training with approximately no positional bias.

#### Frequency expansion

Scalar positions are expanded to *d*_*h*_*/*2 rotation angles at geometric frequencies:

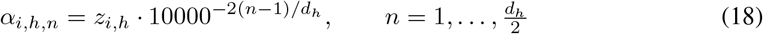

implemented as `z.unsqueeze(-1) * freqs` where `freqs` 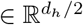.

#### Relative position property

By the RoPE inner product identity, the attention score between genes *i* and *j* satisfies:

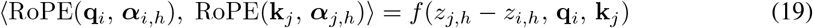

so attention depends only on the relative scalar displacement *z*_*j,h*_− *z*_*i,h*_. Because **h**^(*l*)^ is conditioned on the perturbation identity via the adapter at every layer, *z*_*i,h*_ tracks each gene’s functional role under the specific perturbation at inference time without any explicit supervision on gene-gene relationships.

##### Algorithm 2

Adaptive Repositioning encoding (per attention operation, per layer)

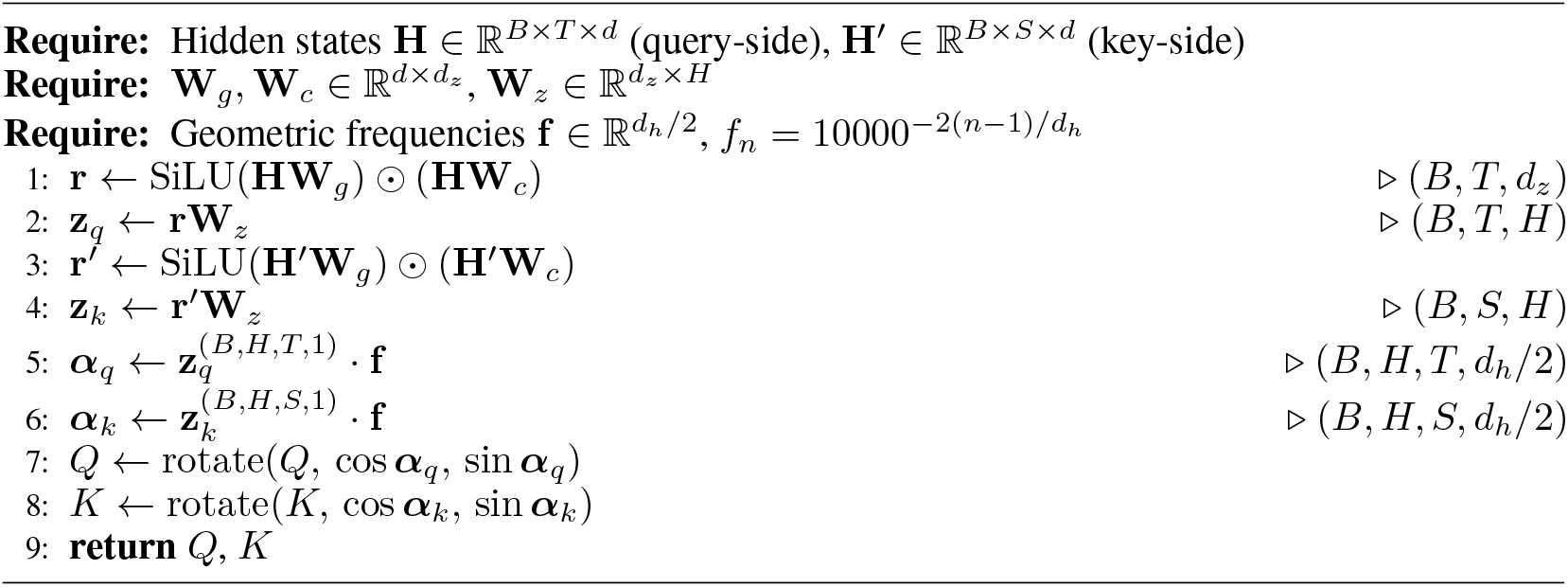

### B.3 Combined Rotation

When both mechanisms are active, the total rotation angle applied at each attention operation is the sum:

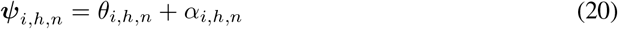

The combined pairwise score between genes *i* and *j* depends on:

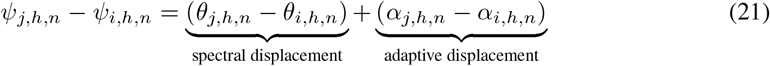

The spectral term encodes fixed co-expression proximity from the Laplacian; the adaptive term encodes perturbation-conditioned functional displacement. Because the angles are summed before the single RoPE rotation, no additional parameters are required to combine them.

## C. Model Architecture and Implementation Details

### Gene Encoder

Gene token ids are mapped to embeddings via `nn.Embedding` and layer-normalized. When spectral positional encoding is enabled, a `GeneAttention` layer applies spectral coexpression rotary encoding over the precomputed Laplacian eigenvectors before the embeddings are consumed by the backbone. The attention mask restricts each query gene to attend only to its top-*k* co-expression neighbors in the gene vocabulary, implemented as a dense additive float mask initialized to −∞ with zeros placed at valid neighbor positions. The gene encoder is shared between the gene identity pathway and the perturbation encoding pathway: knocked-out gene ids are passed through the same encoder and mean-pooled to produce the perturbation embedding **p**.

### Backbone

The backbone is a stack of *L* `DiffPerceiverBlock` layers. Each block consists of two `DifferentialTransformerBlock` passes in sequence. The first is a self-attention pass over the noisy state **Y** against itself; spectral coordinates are disabled here since spectral coexpression encodes inter-gene topology rather than intra-sequence position, though AR still runs and assigns content-based relative positions. The second is a differential contrast pass with queries from **Y** and keys/values from the control cell **X**_ctrl_; both SPE and AR are active here.

Within each `MultiheadDiffAttn`, four independent linear projections produce (*Q*_1_, *K*_1_) from the primary input and (*Q*_2_, *K*_2_) from the secondary input. The differential attention map is:

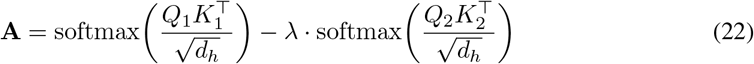

where 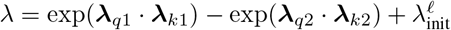 with 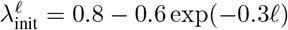. The output is normalized per-head by RMSNorm and scaled by 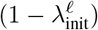.

At every attention operation, queries and keys are rotated by the combined angle ***ψ*** = ***θ*** + ***α***. The spectral coexpression graph component ***θ*** is computed as *θ*_*h,n*_ = Σ_*k*_ Ω_*h,n,k*_ *s*_*i,k*_ via a batched einsum over spectral coordinates and the learnable frequency tensor **Ω**. The AR component ***α*** is computed per layer from the current hidden state through `GeneAdaptiveRepositioning`:

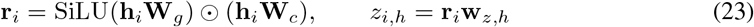

and converted to rotation angles at geometric frequencies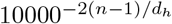 before application.

### Layer conditioning

Before each block, gene identity is injected via `GeneadaLN`, which applies an adaptive layer norm whose shift, scale, and gate are derived from the gene embeddings. The perturbation embedding is then concatenated with the hidden state along the feature dimension and projected back to ℝ^*d*^ via a two-layer LeakyReLU adapter before attention is computed. Timestep conditioning is applied inside each `DifferentialTransformerBlock` via adaLN-Zero modulation: a SiLU-linear network maps the timestep embedding **t**_emb_ ∈ℝ^*d*^ to six modulation scalars (shift, scale, gate for attention; shift, scale, gate for MLP).

### Value encoding

Expression values are encoded by two independent `ContinuousValueEncoder` instances one for the noisy state **x**_*t*_, one for the control cell **c**_ctrl_ each a two-layer ReLU MLP with layer normalization and a clamp at `max_value`. The token representation entering the backbone is the sum of the value embedding and the corresponding gene embedding from the spectral encoder. The two token sequences are fused by concatenation along the feature dimension followed by a two-layer GELU MLP with layer normalization.

### Output head

The final per-gene hidden state is concatenated with the perturbation embedding and passed through a three-layer LeakyReLU MLP (`ExprDecoder`) that projects to a scalar expression prediction per gene. Outputs are clamped to R_≥0_ at inference.

### Initialization

All linear layers use Xavier uniform initialization. **W**_*z*_ in each `GeneAdaptiveRepositioning` module is re-initialized to *N* (0, 0.01^2^) after the global Xavier pass to preserve the near-zero AR start. **Ω** in `Spectral Coexpression Rotary Encoding` is initialized with a near-diagonal structure placing unit weight on the *f* mod *K* eigenvector component for dimension pair *f* .

## D Hyperparameters and Reproduction

### Preprocessing

For each dataset, the top-*K*_HVG_ highly variable genes are selected and log-normalized. The co-expression adjacency matrix is constructed from the training split only. Laplacian eigenvectors are computed once as a preprocessing step and saved as a fixed tensor indexed by vocabulary token id. All experiments were ran on NVidia H100 GPUs with 80GB VRAM.

### Spectral encoding

The eigenvector dimension is *K* = 32. The top-*k* sparsification uses *k* = 30 neighbors per gene with negative edge retention enabled, symmetrizing the adjacency via elementwise max. The resulting spectral coordinate tensor has shape |*V*| × 32 and is frozen throughout training.

### Adaptive repositioning

The bottleneck dimension is *d*_*z*_ = max(*d/*8, 16). **W**_*z*_ is initialized at *σ* = 0.01.

## NeurIPS Paper Checklist

The checklist is designed to encourage best practices for responsible machine learning research, addressing issues of reproducibility, transparency, research ethics, and societal impact. Do not remove the checklist: **The papers not including the checklist will be desk rejected**. The checklist should follow the references and follow the (optional) supplemental material. The checklist does NOT count towards the page limit.

Please read the checklist guidelines carefully for information on how to answer these questions. For each question in the checklist:

- You should answer [Yes], [No], or [N/A].
- [N/A] means either that the question is Not Applicable for that particular paper or the relevant information is Not Available.
- Please provide a short (1–2 sentence) justification right after your answer (even for [N/A]).

### The checklist answers are an integral part of your paper submission

They are visible to the reviewers, area chairs, senior area chairs, and ethics reviewers. You will also be asked to include it (after eventual revisions) with the final version of your paper, and its final version will be published with the paper.

The reviewers of your paper will be asked to use the checklist as one of the factors in their evaluation. While [Yes] is generally preferable to [No], it is perfectly acceptable to answer [No] provided a proper justification is given (e.g., error bars are not reported because it would be too computationally expensive” or “we were unable to find the license for the dataset we used”). In general, answering [No] or [N/A] is not grounds for rejection. While the questions are phrased in a binary way, we acknowledge that the true answer is often more nuanced, so please just use your best judgment and write a justification to elaborate. All supporting evidence can appear either in the main paper or the supplemental material, provided in appendix. If you answer [Yes] to a question, in the justification please point to the section(s) where related material for the question can be found.

IMPORTANT, please:

- **Delete this instruction block, but keep the section heading “NeurIPS Paper Checklist”**,
- **Keep the checklist subsection headings, questions/answers and guidelines below**.
- **Do not modify the questions and only use the provided macros for your answers**.

1. **Claims** Question: Do the main claims made in the abstract and introduction accurately reflect the paper’s contributions and scope? Answer: [Yes] Justification: Yes the title, abstract, and introduction mentions name of our method and application that we show for it along with the contributions and scope. Guidelines:
  - The answer [N/A] means that the abstract and introduction do not include the claims made in the paper.
  - The abstract and/or introduction should clearly state the claims made, including the contributions made in the paper and important assumptions and limitations. A [No] or [N/A] answer to this question will not be perceived well by the reviewers.
  - The claims made should match theoretical and experimental results, and reflect how much the results can be expected to generalize to other settings.
  - It is fine to include aspirational goals as motivation as long as it is clear that these goals are not attained by the paper.
2. **Limitations** Question: Does the paper discuss the limitations of the work performed by the authors? Answer: [Yes] Justification: Yes we have a dedicated limitations section in the end. Guidelines:
  - The answer [N/A] means that the paper has no limitation while the answer [No] means that the paper has limitations, but those are not discussed in the paper.
  - The authors are encouraged to create a separate “Limitations” section in their paper.
  - The paper should point out any strong assumptions and how robust the results are to violations of these assumptions (e.g., independence assumptions, noiseless settings, model well-specification, asymptotic approximations only holding locally). The authors should reflect on how these assumptions might be violated in practice and what the implications would be.
  - The authors should reflect on the scope of the claims made, e.g., if the approach was only tested on a few datasets or with a few runs. In general, empirical results often depend on implicit assumptions, which should be articulated.
  - The authors should reflect on the factors that influence the performance of the approach. For example, a facial recognition algorithm may perform poorly when image resolution is low or images are taken in low lighting. Or a speech-to-text system might not be used reliably to provide closed captions for online lectures because it fails to handle technical jargon.
  - The authors should discuss the computational efficiency of the proposed algorithms and how they scale with dataset size.
  - If applicable, the authors should discuss possible limitations of their approach to address problems of privacy and fairness.
  - While the authors might fear that complete honesty about limitations might be used by reviewers as grounds for rejection, a worse outcome might be that reviewers discover limitations that aren’t acknowledged in the paper. The authors should use their best judgment and recognize that individual actions in favor of transparency play an important role in developing norms that preserve the integrity of the community. Reviewers will be specifically instructed to not penalize honesty concerning limitations.
3. **Theory assumptions and proofs** Question: For each theoretical result, does the paper provide the full set of assumptions and a complete (and correct) proof? Answer: [Yes] Justification: Yes all assumptions and proofs with images, analysis and metrics are provided for our results Guidelines:
  - The answer [N/A] means that the paper does not include theoretical results.
  - All the theorems, formulas, and proofs in the paper should be numbered and cross-referenced.
  - All assumptions should be clearly stated or referenced in the statement of any theorems.
  - The proofs can either appear in the main paper or the supplemental material, but if they appear in the supplemental material, the authors are encouraged to provide a short proof sketch to provide intuition.
  - Inversely, any informal proof provided in the core of the paper should be complemented by formal proofs provided in appendix or supplemental material.
  - Theorems and Lemmas that the proof relies upon should be properly referenced.
4. **Experimental result reproducibility** Question: Does the paper fully disclose all the information needed to reproduce the main experimental results of the paper to the extent that it affects the main claims and/or conclusions of the paper (regardless of whether the code and data are provided or not)? Answer: [Yes] Justification: Yes the paper provides the data split, algorithms, architecture, and hyperparameter details to reproduce the experiments. Guidelines:
  - The answer [N/A] means that the paper does not include experiments.
  - If the paper includes experiments, a [No] answer to this question will not be perceived well by the reviewers: Making the paper reproducible is important, regardless of whether the code and data are provided or not.
  - If the contribution is a dataset and/or model, the authors should describe the steps taken to make their results reproducible or verifiable.
  - Depending on the contribution, reproducibility can be accomplished in various ways. For example, if the contribution is a novel architecture, describing the architecture fully might suffice, or if the contribution is a specific model and empirical evaluation, it may be necessary to either make it possible for others to replicate the model with the same dataset, or provide access to the model. In general. releasing code and data is often one good way to accomplish this, but reproducibility can also be provided via detailed instructions for how to replicate the results, access to a hosted model (e.g., in the case of a large language model), releasing of a model checkpoint, or other means that are appropriate to the research performed.
  - While NeurIPS does not require releasing code, the conference does require all submissions to provide some reasonable avenue for reproducibility, which may depend on the nature of the contribution. For example
    a. If the contribution is primarily a new algorithm, the paper should make it clear how to reproduce that algorithm.
    b. If the contribution is primarily a new model architecture, the paper should describe the architecture clearly and fully.
    c. If the contribution is a new model (e.g., a large language model), then there should either be a way to access this model for reproducing the results or a way to reproduce the model (e.g., with an open-source dataset or instructions for how to construct the dataset).
    d. We recognize that reproducibility may be tricky in some cases, in which case authors are welcome to describe the particular way they provide for reproducibility. In the case of closed-source models, it may be that access to the model is limited in some way (e.g., to registered users), but it should be possible for other researchers to have some path to reproducing or verifying the results.
5. **Open access to data and code** Question: Does the paper provide open access to the data and code, with sufficient instructions to faithfully reproduce the main experimental results, as described in supplemental material? Answer: [Yes] Justification: Yes the paper provides full code as supplementary material along with links to the preprocessed data to reprodice all experiments. Guidelines:
  - The answer [N/A] means that paper does not include experiments requiring code.
  - Please see the NeurIPS code and data submission guidelines (https://neurips.cc/public/guides/CodeSubmissionPolicy) for more details.
  - While we encourage the release of code and data, we understand that this might not be possible, so [No] is an acceptable answer. Papers cannot be rejected simply for not including code, unless this is central to the contribution (e.g., for a new open-source benchmark).
  - The instructions should contain the exact command and environment needed to run to reproduce the results. See the NeurIPS code and data submission guidelines (https://neurips.cc/public/guides/CodeSubmissionPolicy) for more details.
  - The authors should provide instructions on data access and preparation, including how to access the raw data, preprocessed data, intermediate data, and generated data, etc.
  - The authors should provide scripts to reproduce all experimental results for the new proposed method and baselines. If only a subset of experiments are reproducible, they should state which ones are omitted from the script and why.
  - At submission time, to preserve anonymity, the authors should release anonymized versions (if applicable).
  - Providing as much information as possible in supplemental material (appended to the paper) is recommended, but including URLs to data and code is permitted.
6. **Experimental setting/details** Question: Does the paper specify all the training and test details (e.g., data splits, hyperparameters, how they were chosen, type of optimizer) necessary to understand the results? Answer: [Yes] Justification: Yes the GPU VRAM memory, data slits, hyperparameter values, and optimzer details are mentioned in the appendix. Guidelines:
  - The answer [N/A] means that the paper does not include experiments.
  - The experimental setting should be presented in the core of the paper to a level of detail that is necessary to appreciate the results and make sense of them.
  - The full details can be provided either with the code, in appendix, or as supplemental material.
7. **Experiment statistical significance** Question: Does the paper report error bars suitably and correctly defined or other appropriate information about the statistical significance of the experiments? Answer: [Yes] Justification: Yes the paper reports error bars for the biological validation results along with p values using paired wilcoxon tests, and single fixed seed results for ablation studies and training curves. Guidelines:
  - The answer [N/A] means that the paper does not include experiments.
  - The authors should answer [Yes] if the results are accompanied by error bars, confidence intervals, or statistical significance tests, at least for the experiments that support the main claims of the paper.
  - The factors of variability that the error bars are capturing should be clearly stated (for example, train/test split, initialization, random drawing of some parameter, or overall run with given experimental conditions).
  - The method for calculating the error bars should be explained (closed form formula, call to a library function, bootstrap, etc.)
  - The assumptions made should be given (e.g., Normally distributed errors).
  - It should be clear whether the error bar is the standard deviation or the standard error of the mean.
  - It is OK to report 1-sigma error bars, but one should state it. The authors should preferably report a 2-sigma error bar than state that they have a 96% CI, if the hypothesis of Normality of errors is not verified.
  - For asymmetric distributions, the authors should be careful not to show in tables or figures symmetric error bars that would yield results that are out of range (e.g., negative error rates).
  - If error bars are reported in tables or plots, the authors should explain in the text how they were calculated and reference the corresponding figures or tables in the text.
8. **Experiments compute resources** Question: For each experiment, does the paper provide sufficient information on the computer resources (type of compute workers, memory, time of execution) needed to reproduce the experiments? Answer: [Yes] Justification: Yes the compute resource details are provided in the appendix. Guidelines:
  - The answer [N/A] means that the paper does not include experiments.
  - The paper should indicate the type of compute workers CPU or GPU, internal cluster, or cloud provider, including relevant memory and storage.
  - The paper should provide the amount of compute required for each of the individual experimental runs as well as estimate the total compute.
  - The paper should disclose whether the full research project required more compute than the experiments reported in the paper (e.g., preliminary or failed experiments that didn’t make it into the paper).
9. **Code of ethics** Question: Does the research conducted in the paper conform, in every respect, with the NeurIPS Code of Ethics https://neurips.cc/public/EthicsGuidelines? Answer: [Yes] Justification: Yes code of ethics is strictly followed. Guidelines:
  - The answer [N/A] means that the authors have not reviewed the NeurIPS Code of Ethics.
  - If the authors answer [No], they should explain the special circumstances that require a deviation from the Code of Ethics.
  - The authors should make sure to preserve anonymity (e.g., if there is a special consideration due to laws or regulations in their jurisdiction).
10. **Broader impacts** Question: Does the paper discuss both potential positive societal impacts and negative societal impacts of the work performed? Answer: [N/A] Justification: Our work is based on encoding methods for cellular perturbation modeling and does not include any societal impact studies. Guidelines:
  - The answer [N/A] means that there is no societal impact of the work performed.
  - If the authors answer [N/A] or [No], they should explain why their work has no societal impact or why the paper does not address societal impact.
  - Examples of negative societal impacts include potential malicious or unintended uses (e.g., disinformation, generating fake profiles, surveillance), fairness considerations (e.g., deployment of technologies that could make decisions that unfairly impact specific groups), privacy considerations, and security considerations.
  - The conference expects that many papers will be foundational research and not tied to particular applications, let alone deployments. However, if there is a direct path to any negative applications, the authors should point it out. For example, it is legitimate to point out that an improvement in the quality of generative models could be used to generate Deepfakes for disinformation. On the other hand, it is not needed to point out that a generic algorithm for optimizing neural networks could enable people to train models that generate Deepfakes faster.
  - The authors should consider possible harms that could arise when the technology is being used as intended and functioning correctly, harms that could arise when the technology is being used as intended but gives incorrect results, and harms following from (intentional or unintentional) misuse of the technology.
  - If there are negative societal impacts, the authors could also discuss possible mitigation strategies (e.g., gated release of models, providing defenses in addition to attacks, mechanisms for monitoring misuse, mechanisms to monitor how a system learns from feedback over time, improving the efficiency and accessibility of ML).
11. **Safeguards** Question: Does the paper describe safeguards that have been put in place for responsible release of data or models that have a high risk for misuse (e.g., pre-trained language models, image generators, or scraped datasets)? Answer: [N/A] Justification: Our work is based on encoding methods for cellular perturbation modeling and does not include any safeguard or risk analysis studies. Guidelines:
  - The answer [N/A] means that the paper poses no such risks.
  - Released models that have a high risk for misuse or dual-use should be released with necessary safeguards to allow for controlled use of the model, for example by requiring that users adhere to usage guidelines or restrictions to access the model or implementing safety filters.
  - Datasets that have been scraped from the Internet could pose safety risks. The authors should describe how they avoided releasing unsafe images.
  - We recognize that providing effective safeguards is challenging, and many papers do not require this, but we encourage authors to take this into account and make a best faith effort.
12. **Licenses for existing assets** Question: Are the creators or original owners of assets (e.g., code, data, models), used in the paper, properly credited and are the license and terms of use explicitly mentioned and properly respected? Answer: [Yes] Justification: Yes all previous methods we compare against and datasets we evaluate on have been cited. Guidelines:
  - The answer [N/A] means that the paper does not use existing assets.
  - The authors should cite the original paper that produced the code package or dataset.
  - The authors should state which version of the asset is used and, if possible, include a URL.
  - The name of the license (e.g., CC-BY 4.0) should be included for each asset.
  - For scraped data from a particular source (e.g., website), the copyright and terms of service of that source should be provided.
  - If assets are released, the license, copyright information, and terms of use in the package should be provided. For popular datasets, paperswithcode.com/datasets has curated licenses for some datasets. Their licensing guide can help determine the license of a dataset.
  - For existing datasets that are re-packaged, both the original license and the license of the derived asset (if it has changed) should be provided.
  - If this information is not available online, the authors are encouraged to reach out to the asset’s creators.
13. **New assets** Question: Are new assets introduced in the paper well documented and is the documentation provided alongside the assets? Answer: [Yes] Justification: Yes new encoding method has been well documented and code is provided as supplementary Guidelines:
  - The answer [N/A] means that the paper does not release new assets.
  - Researchers should communicate the details of the dataset/code/model as part of their submissions via structured templates. This includes details about training, license, limitations, etc.
  - The paper should discuss whether and how consent was obtained from people whose asset is used.
  - At submission time, remember to anonymize your assets (if applicable). You can either create an anonymized URL or include an anonymized zip file.
14. **Crowdsourcing and research with human subjects** Question: For crowdsourcing experiments and research with human subjects, does the paper include the full text of instructions given to participants and screenshots, if applicable, as well as details about compensation (if any)? Answer: [N/A] Justification: No crowdsourcing experiments were performed. Guidelines:
  - The answer [N/A] means that the paper does not involve crowdsourcing nor research with human subjects.
  - Including this information in the supplemental material is fine, but if the main contribution of the paper involves human subjects, then as much detail as possible should be included in the main paper.
  - According to the NeurIPS Code of Ethics, workers involved in data collection, curation, or other labor should be paid at least the minimum wage in the country of the data collector.
15. **Institutional review board (IRB) approvals or equivalent for research with human subjects** Question: Does the paper describe potential risks incurred by study participants, whether such risks were disclosed to the subjects, and whether Institutional Review Board (IRB) approvals (or an equivalent approval/review based on the requirements of your country or institution) were obtained? Answer: [N/A] Justification: Our work does not include any human subjects Guidelines:
  - The answer [N/A] means that the paper does not involve crowdsourcing nor research with human subjects.
  - Depending on the country in which research is conducted, IRB approval (or equivalent) may be required for any human subjects research. If you obtained IRB approval, you should clearly state this in the paper.
  - We recognize that the procedures for this may vary significantly between institutions and locations, and we expect authors to adhere to the NeurIPS Code of Ethics and the guidelines for their institution.
  - For initial submissions, do not include any information that would break anonymity (if applicable), such as the institution conducting the review.
16. **Declaration of LLM usage** Question: Does the paper describe the usage of LLMs if it is an important, original, or non-standard component of the core methods in this research? Note that if the LLM is used only for writing, editing, or formatting purposes and does *not* impact the core methodology, scientific rigor, or originality of the research, declaration is not required. Answer: [N/A] Justification: Our work did not use LLM in any way for development of methods. LLMs were only used to polish text writing and grammar. Guidelines:
  - The answer [N/A] means that the core method development in this research does not involve LLMs as any important, original, or non-standard components.
  - Please refer to our LLM policy in the NeurIPS handbook for what should or should not be described.

